# “Less is more”: a dose-response account of intranasal oxytocin pharmacodynamics in the human brain

**DOI:** 10.1101/2021.01.18.427062

**Authors:** Daniel Martins, Katja Broadmann, Mattia Veronese, Ottavia Dipasquale, Ndaba Mazibuko, Uwe Schuschnig, Fernando Zelaya, Aikaterini Fotopoulou, Yannis Paloyelis

## Abstract

Intranasal oxytocin is receiving increasing research attention as a potential treatment for several brain disorders due to promising preclinical results. However, translating these findings to humans has been hampered by remaining uncertainties about its pharmacodynamics and the methods used to probe its effects in the human brain. Using a dose-response design (9, 18 and 36 IU) and a novel administration device, we demonstrate that oxytocin-induced physiological changes in the amygdala at rest, a key-hub of the brain oxytocin system, follow a dose-response curve with maximal effects for lower doses. Yet, the effects of oxytocin vary by amygdala subdivision, highlighting the need to qualify dose-response curves within subregion. We further link physiological changes with the density of the oxytocin receptor gene mRNA across brain regions, strengthening our confidence in intranasal oxytocin as a valid approach to engage central targets. Finally, we demonstrate that intranasal oxytocin does not disrupt cerebrovascular reactivity; the absence of major vascular confounds corroborates the validity of haemodynamic neuroimaging to probe the effects of intranasal oxytocin in the human brain.

## Introduction

Intranasal oxytocin, the most widely used method for oxytocin administration in humans, has been suggested as a promising therapeutic strategy for several brain disorders where we currently lack effective treatments (e.g., autism spectrum disorder^1^, schizophrenia^2^, migraine^3^, stroke^4^, obesity^5^, Prader-Willi^6^). An increasing number of clinical trials have been evaluating the efficacy of specific nominal doses of intranasal oxytocin (for an overview see^7–11^), yet most studies have been inconclusive at best^11, 12^. The lack of unequivocal findings regarding the effects of intranasal oxytocin in trials with human patients contrasts with a flood of evidence showing consistent beneficial effects in animal models^13, 14^, where oxytocin is often administered centrally^15–17^. This discrepancy has raised questions around the basic pharmacology of intranasal oxytocin, and regarding the neuroimaging measures used to assess its effects on the human brain, namely: (i) which doses might be most effective in targeting specific brain circuits^18, 19^? (ii) Do the changes in brain physiology that follow the administration of intranasal oxytocin result from the engagement of the OXTR, the primary oxytocin target?? (iii) Could the hemodynamic neuroimaging markers typically used to probe the effects of intranasal oxytocin in the human brain be affected by major unspecific vascular confounds?

Regarding the first question, the most commonly used dose in studies administering intranasal oxytocin with nasal sprays (24IU) has been selected largely due to historical precedence^20^, rather than a systematic investigation of dose-response curves for each targeted brain region^21, 22^. However, as with many drugs, dose is likely to play an important role in determining the response to intranasal oxytocin. First, when binding the oxytocin receptor (OXTR)^23^, oxytocin recruits different intracellular G protein pathways with opposite effects on neuronal activity, depending on the amount that is available extracellularly^24, 25^. Increases in oxytocin bioavailability shift OXTR-coupling away from the excitatory G_q_ to the inhibitory G_i_ protein^24, 25^. Hence, as oxytocin bioavailability increases, the recruitment of the inhibitory G_i_ pathway is likely to counteract the effects related to the recruitment of the excitatory G_q_ pathway, resulting in null or even opposing drug effects^24, 25^. Second, increases in local oxytocin bioavailability also increase the chance of engagement of vasopressin receptors, such as the V1aR, for which oxytocin has lower affinity than the OXTR^26^. This cross-receptor activation may ultimately counteract the effects related to OXTR engagement^27^. Ultimately, the complexity associated with oxytocin signalling described above makes it difficult to predict what the effects of specific nominal doses of intranasal oxytocin might be on brain circuits that vary in the availability of oxytocin targets, without detailed region-specific characterizations of dose-response.

Previous studies have reported dose-dependent effects of oxytocin in both non-human^28, 29^ and human^30–35^ species. This evidence was gathered in blood oxygen level-dependent (BOLD)-functional magnetic resonance imaging (fMRI) task-based studies, using paradigms that engage specific neural circuits, such as the amygdala^30, 32, 33^, or measuring specific neurophysiological processes, such as pupil reactivity to emotional faces^32^ or chemosensory decoding^35^. Overall, these studies suggested that lower-to-medium doses (8-24 IU) of oxytocin may be more efficacious than higher doses^36^. However, how well these dose-response effects might extend beyond the specific processes engaged by the tasks employed is currently unclear. Moreover, the investigation of pharmacological effects using BOLD fMRI, which is a relative measure involving the comparison of two conditions, cannot disentangle the modulatory effects of the drug on task-dependent activation states from changes in baseline brain function^37^.

To overcome these limitations, in this study we investigate dose-related changes in local perfusion by measuring regional cerebral blood flow (rCBF) at rest, which allows us to uncover basic pharmacological mechanisms that are not restricted to the circuits engaged by specific paradigms^38^. We have demonstrated the sensitivity of arterial spin labelling (ASL) MRI in quantifying changes in regional cerebral blood flow (rCBF) at rest after intranasal oxytocin administration^22, 39^. Changes in rCBF at rest provide a quantitative, non-invasive pharmacodynamic marker of the effects of acute doses of psychoactive drugs^40, 41^, with high spatial resolution and excellent temporal reproducibility^42^. As a result of neuro-vascular coupling, changes in rCBF in response to oxytocin are likely to reflect changes in metabolic demand associated with neuronal activity^43^.

We focused on dose-related changes in local rCBF in a key-hub of the brain oxytocin system, the amygdala. Modulation of amygdala’s activity constitutes one of the most robust findings in animal studies and intranasal oxytocin studies in humans^44–48^, rendering it a well justified target. However, studies in rodents^27, 48, 49^ and humans^50^ have provided evidence that the effects of oxytocin vary by amygdala subdivision. Yet how each subdivision responds to different doses of intranasal oxytocin is currently unknown, highlighting the need to disentangle the complex modulatory role of oxytocin on different amygdalar circuits. Therefore, in this study, we also conducted secondary analyses to examine separately the four main amygdala subdivisions, centromedial, laterobasal, superficial and amygdalostriatal transition area, to characterize dose-response within each subdivision. In addition to changes in local rCBF, we also assessed functional connectivity by employing a new approach based on group-based rCBF covariance statistics^51^. This approach has been shown to have exquisite sensitivity to detect brain-wide effects of neuromodulators^52–55^, including intranasal oxytocin in rodents^56^; however, it has never been used before to investigate the effects of intranasal oxytocin on brain physiology in humans.

Regarding the second question, it is currently unclear whether the physiological brain changes (i.e. rCBF) that follow the administration of intranasal oxytocin do result from the engagement of the OXTR, the primary oxytocin target. While recent studies in primates^57^ and rodents^58^ have shown that synthetic oxytocin when administered intranasally can reach the brain parenchyma, linking the functional effects of intranasal oxytocin in the brain to the engagement of its receptor would be critical to strengthening our confidence in the validity of using intranasal oxytocin as a method to target the central oxytocin system. This question could be neatly addressed by investigating whether a brain-penetrant, specific OXTR antagonist could blunt the changes in rCBF induced by intranasal oxytocin. Nevertheless, such an antagonist is not currently available for use in humans. Therefore, in this study we addressed this question indirectly, by combining our neuroimaging data with transcriptomic data from the Allen Brain Atlas^59^ to investigate if the distribution of the levels of mRNA of the *OXTR* gene in the post-mortem human brain can predict the rCBF changes that follow the administration of the three doses of intranasal oxytocin that we employed.

Regarding the third question, haemodynamic neuroimaging measures are widely used as a probe^60^ of the effects of intranasal oxytocin on brain function, however the validity of this approach remains to be confirmed. Changes in haemodynamic MRI measures reflect a complex cascade of cellular, metabolic and vascular events associated with changes in neuronal activity^38^. The main effects of drugs on brain’s physiology, such as the effects on rCBF we have reported for intranasal oxytocin in our previous studies^22, 39^, are typically interpreted as the result of enhanced or decreased pre-or post-synaptic activity due to the action of the drug on its targets^38^. This approach is founded on the assumption that oxytocin does not interfere with the ability of the cerebral vasculature to modulate blood flow in response to vasoactive stimuli such as CO_2_ (cerebrovascular reactivity (CVR)), which could represent a major confound. For instance, an undetected drug-induced modulation of CVR could produce a BOLD response in the absence of a change in neural activity, or “mask” an actual neuronal response. This issue is particularly relevant when examining intranasal oxytocin since oxytocin is known to have vasoactive properties^61^. If oxytocin disrupts CVR, then changes in MRI hemodynamic measures, such as changes in the BOLD signal, cannot be directly attributed to the modulation of neuronal activity. Hence, following several recent recommendations to address the potential impact that pharmacological compounds may have on the cerebrovasculature^62–64^, we investigated the presence of unspecific effects of intranasal oxytocin on CVR during a breath hold task^65^.

To investigate these three main questions, we employed a double-blind, placebo-controlled, crossover design where we administered three different doses of intranasal oxytocin (9IU, 18IU and 36IU) or placebo to 24 healthy men using a novel administration device (PARI SINUS nebuliser) that maximizes deposition in areas of putative nose-to-brain transport (see Figure 1 for a summary of our experimental protocol). We demonstrate that intranasal oxytocin-induced changes in rCBF in the amygdala and rCBF covariance with other key hubs of the central oxytocin system follow a dose-response curve with lower doses inducing the highest changes from placebo. However, we also demonstrate that one of the amygdala subdivision responds to oxytocin with a different dose-response profile than the remaining subdivisions, which highlights the need to qualify the selection of dose for maximal pharmacologic effects within subdivision. We provide indirect evidence supporting the validity of using intranasal oxytocin as a pharmacological method to target the brain’s oxytocin system by showing that the rCBF changes induced by the effective doses can be predicted by the expression of the mRNA of *OXTR* across regions of the whole-brain. Furthermore, we also confirm the validity of neuroimaging measures to investigate the effects of intranasal oxytocin on human brain function by demonstrating that none of the doses of oxytocin tested here affect CVR, hence confirming the absence of major vascular confounds across a range of doses.

**Fig. 1.**
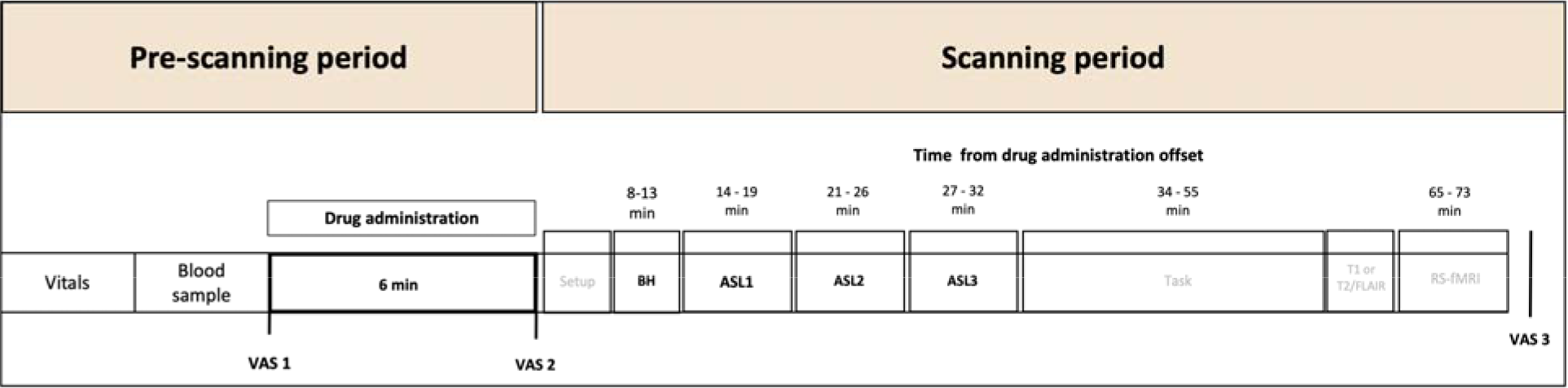
Overview of the experimental protocol. In this diagram, we provide an overview of the experimental procedures of our study. <u>Pre-Scanning period:</u> Each session started with a quick assessment of vitals (heart rate and blood pressure) and collection of two blood samples for plasma isolation. Then, participants self-administered one of three possible doses of intranasal oxytocin (∼9, 18 or 36 IU) or placebo using the PARI SINUS nebulizer. The participants used the nebulizer for 3 mins in each nostril (total administration 6 mins). Immediately before and after drug administration, participants filled a battery of visual analog scales (VAS) to assess subjective drug effects (alertness, mood and anxiety). <u>Scanning Period:</u> Participants were then guided to a magnetic resonance imaging scanner, where acquired BOLD-fMRI during a breath hold task, three consecutive arterial spin labelling scans, one BOLD-fMRI prosocial learning task, followed by structural scans and one BOLD-fMRI resting-state at the end. We present the time-interval post-dosing (mean time from drug administration offset) during which each scan took place. At the end of the scanning session, we repeated the same battery of VAS to subjective drug effects.

## Results

### Dose-response effects of intranasal oxytocin on local rCBF

To answer our first question, we used ASL to investigate how different doses of intranasal oxytocin impact on local rCBF and functional connectivity of the amygdala and its subdivisions at rest at about 14-32 mins post-dosing (Figure 1). First, we conducted some control analyses to investigate treatment (placebo, low, medium or high doses of intranasal oxytocin), time-interval (14-19 min, 21-26 min and 27-32 min post-dosing) and treatment x time-interval effects on subjective drug effects (alertness, mood and anxiety) and global CBF. Treatment did not impact on alertness, mood or anxiety (see Supplementary Table 1 and Supplementary Figure 1 for further details). On global CBF, only the main effect of treatment was significant (F(3,105.505) = 4.666, p = 0.004) (Supplementary Table 2). Compared to placebo, all three doses decreased global CBF (see Supplementary Figure 2 for further details). The existence of significant treatment effects on global CBF supported our rationale for including global CBF as a confounding covariate in all of our rCBF analyses, consistent with previous standard practice^39^.

We followed up the control analyses by investigating treatment, time-interval and treatment x time-interval effects on extracted values of rCBF at rest focusing on our selected ROIs: i) the right and left amygdala (whole), as primary outcomes; and ii) their respective centromedial, laterobasal, superficial and amygdalostriatal transition area subdivisions, as secondary outcomes. We also conducted exploratory whole-brain analyses to investigate other potential treatment, time-interval and treatment x time-interval effects on rCBF at rest beyond the amygdala. We describe the results of each of these analyses below.

#### Amygdala (whole)

We found a significant main effect of treatment for the left amygdala (Table 1). None of the post-hoc comparisons with placebo or between active doses survived correction (smallest p_adjusted_ = 0.072). Numerically, the low dose produced the largest nominal decrease in rCBF compared to placebo (d = 0.213) among the three doses, followed by the medium (d=0.205) and the high dose (d = 0.110). The effects of time-interval or treatment x time-interval were not significant.

**Table 1.** Effects of treatment, time-interval and treatment x time-interval on rCBF in the amygdala and its subdivisions. We investigated the effects of treatment, time-interval and treatment x time-interval on rCBF for 10 anatomical regions-of-interest (ROIs) including the right and left amygdala (whole) and its respective centromedial, laterobasal, superficial and amygdalostriatal transition area subdivisions (left panel). We used a linear mixed-model, considering treatment and time-interval as fixed factors, with a random intercept for subjects and global CBF as a covariate. *p-FDR* values reflect adjusted p-values for the total number of ROIs tested (n=10), using the Benjamini-Hochberg procedure. Statistical significance was set at *p* < 0.05 (two-tailed).

| Region-of-interest | Main effect of treatment |  |  | Main effect of time |  |  | Interaction<br>Treatment x Time |  |  |
| --- | --- | --- | --- | --- | --- | --- | --- | --- | --- |
|  | F | p-value | p-FDR* | F | P-value | p-FDR* | F | p-value | p-FDR* |
| Right amygdala | F(3,99.506) = 0.931 | 0.429 | 0.536 | F(2,168.916) = 2.416 | 0.092 | 0.307 | F(6,95.746) = 0.547 | 0.771 | 1.101 |
| Right Amygdalostriatal Transition Area | F(3,119.772) = 2.712 | 0.048 | 0.080 | F(2,182.374) = 0.657 | 0.520 | 0.867 | F(6,91.028) = 0.770 | 0.595 | 1.488 |
| Right Centromedial amygdala | F(3,132.326) = 1.388 | 0.249 | 0.356 | F(2,158.815) = 0.900 | 0.409 | 0.818 | F(6,94.320) = 0.914 | 0.489 | 1.630 |
| Right Laterobasal amygdala | F(3,132.178) = 0.578 | 0.578 | 0.642 | F(2,155.548) = 0.080 | 0.923 | 0.923 | F(6,99.411) = 0.281 | 0.945 | 0.945 |
| Right Superficial amygdala | F(3,111.448) = 3.753 | 0.013 | 0.043 | F(2,178.391) = 0.293 | 0.746 | 0.933 | F(6,87.407) = 0.451 | 0.843 | 1.054 |
| Left amygdala | F(3,136.397) = 3.326 | 0.022 | 0.044 | F(2,151.584) = 0.348 | 0.707 | 1.010 | F(6,96.587) = 0.337 | 0.916 | 1.018 |
| Left Amygdalostriatal Transition Area | F(3,105.851) = 11.370 | 2.000x10<sup>-6</sup> | 2.000x10<sup>-4</sup> | F(2,166.679) = 3.184 | 0.050 | 0.500 | F(6,84.877) = 0.587 | 0.770 | 1.283 |
| Left Centromedial amygdala | F(3,109.463) = 3.418 | 0.020 | 0.050 | F(2,167.861) = 1.772 | 0.173 | 0.433 | F(6,104.307) = 1.109 | 0.362 | 3.620 |
| Left Laterobasal amygdala | F(3,112.230) = 0.489 | 0.691 | 0.691 | F(2,172.227) = 2.494 | 0.086 | 0.430 | F(6,101.265) = 0.639 | 0.699 | 1.398 |
| Left Superficial amygdala | F(3,128.334) = 6.371 | 4.650x10<sup>-4</sup> | 0.002 | F(2,182.500) = 0.257 | 0.774 | 0.860 | F(6,96.155) = 0.934 | 0.475 | 2.375 |

#### Amygdala (subdivisions)

The effects of time-interval or treatment x time-interval were not significant for any of the amygdala ROIs we tested (Table 1). However, we found significant main effects of treatment for the left centromedial amygdala, the left and right amygdalostriatal transition areas, and the left and right superficial amygdala subdivisions. Only the treatment effects for the right amygdalostriatal transition area ROI did not survive correction for the total number of ROIs tested (Table 1).

For the left centromedial amygdala, the main effect of treatment was driven by decreases in rCBF, compared to placebo, for the low (p_adjusted_ = 0.005) and medium (p_adjusted_ = 0.021) doses but not the high dose (p=0.390). This decrease from placebo was maximal for the low dose (d = 0.261), followed by the medium dose (d = 0.233). Direct comparisons between all possible pairs of our three active doses yielded non-significant differences (smallest p_adjusted_ = 0.058) (Figure 2). For the left amygdalostriatal transition area, only the low dose was significantly lower than placebo (d = 0.859, p_adjusted_ < 0.001) (medium or high dose versus placebo: p_adjusted_ > 0.111). In this ROI, rCBF in the low dose group was also significantly lower than in the medium (p_adjusted_ = 0.002) and high (p_adjusted_ < 0.001) doses groups. Direct comparisons between medium and high doses yielded non-significant differences (p_adjusted_ = 0.595) (Figure 2).

**Fig. 2.**
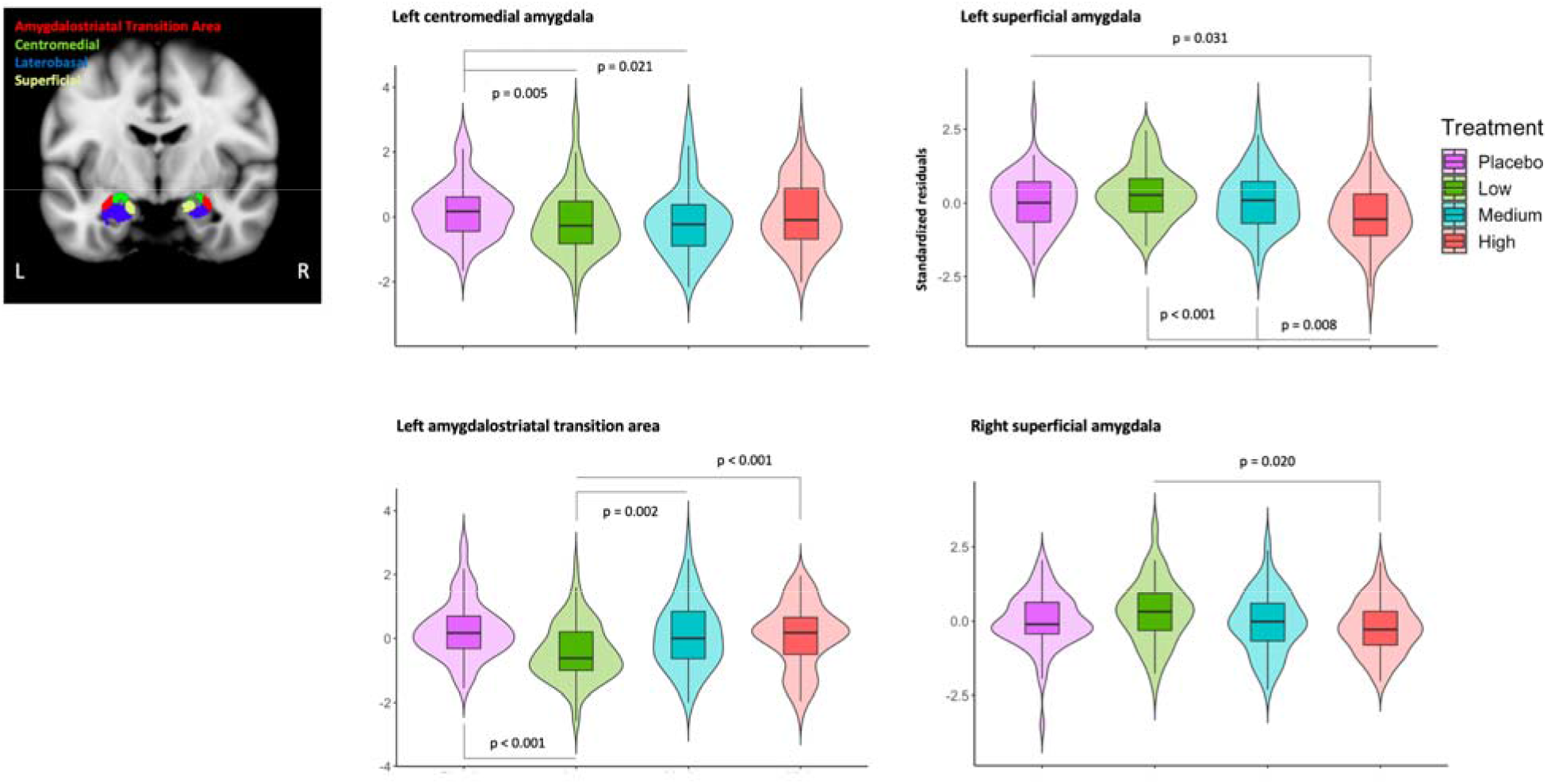
Dose-response effects of intranasal oxytocin on rCBF in the amygdala and its subdivisions. We investigated the effects of treatment, time-interval and treatment x time-interval on rCBF for 10 regions-of-interest (ROIs) including the right and left amygdala and its respective centromedial, laterobasal, superficial and amygdalostriatal transition area subdivisions (left panel), while controlling for global CBF. In the right panel, we present the results of the *post-hoc* investigations of the simple dose effects for the ROIs where we detected a significant main effect of treatment. Statistical significance was set at *p* < 0.05, after correcting for multiple comparisons using the *Sidak* correction.

For the left superficial amygdala ROI, the main effect of treatment was driven by decreases in rCBF, compared to placebo, only for the high dose (p_adjusted_ = 0.031, d=0.526) (Low versus Placebo: p_adjusted_ = 0.123, d = 0.158; Medium versus Placebo: p_adjusted_ = 0.528, d = 0.431). rCBF in left superficial amygdala for the high dose group was also significantly lower than in the low (p_adjusted_ < 0.001) and medium (p_adjusted_ = 0.008) doses groups. Direct comparisons between the low and medium doses yielded non-significant differences (p_adjusted_ = 0.120) (Figure 2). We found a similar pattern of effects for the right superficial amygdala. However, in this case the main effect of treatment was driven by a significant decrease in rCBF after the high dose when compared to the low dose (d = 0.259, p_adjusted_ = 0.020). We did not find any significant differences when we directly compared the high dose against placebo (p_adjusted_ = 0.237, d = 0.434), the low dose against the medium dose (p_adjusted_ = 0.121), or the medium dose against the high dose (p_adjusted_ = 0.437) (Figure 2).

#### Exploratory whole-brain analysis

We conducted exploratory whole-brain analyses to investigate other potential treatment, time-interval and treatment x time-interval effects on rCBF at rest beyond the amygdala. We did not find any cluster depicting significant treatment or time-interval x treatment effects. However, we found three clusters spanning mostly the insula bilaterally that showed a main effect of time-interval (see Supplementary Figure 3).

### Dose-response effects of intranasal oxytocin on functional connectivity using group-based rCBF covariance

In addition to investigating dose-response effects of intranasal oxytocin on local rCBF changes, we examined dose-response effects on functional connectivity within a network of key brain areas of the central oxytocinergic circuitry using group-based rCBF covariance and graph-theory network analysis. Following our analytic strategy on local rCBF, we focused our main analysis on the functional connectivity of the amygdala and its subdivisions with other brain regions considered to be part of the central oxytocinergic circuits, but also conducted exploratory analyses for dose-response changes in the global functional connectivity of our oxytocinergic network as a whole. We describe the results of each of these analyses below.

#### Amygdala (whole)

We found that, compared to placebo, the low dose increased the clustering coefficient of the left amygdala within our oxytocinergic network (p_permuted_ = 0.006) (Figure 3A). Similar comparisons for the medium (p_permuted_ = 0.126) and high (p_permuted_ = 0.112) doses yielded no significant changes, compared to placebo. Direct comparisons between the low and medium, low and high or medium and high doses were not significant (smallest p_permuted_ = 0.123). For node strength, we found a trend for increased node strength of the left amygdala after the low dose (p_permuted_ = 0.066), compared to placebo. Similar comparisons for the medium (p_permuted_ = 0.169) and high (p_permuted_ = 0.739) doses yielded no significant changes. Direct comparisons between the low and medium, low and high or medium and high doses were not significant (smallest p_permuted_ = 0.247). No significant effects on clustering coefficient or node strength were identified for the right amygdala (smallest p_permuted_ = 0.090).

**Fig. 3.**
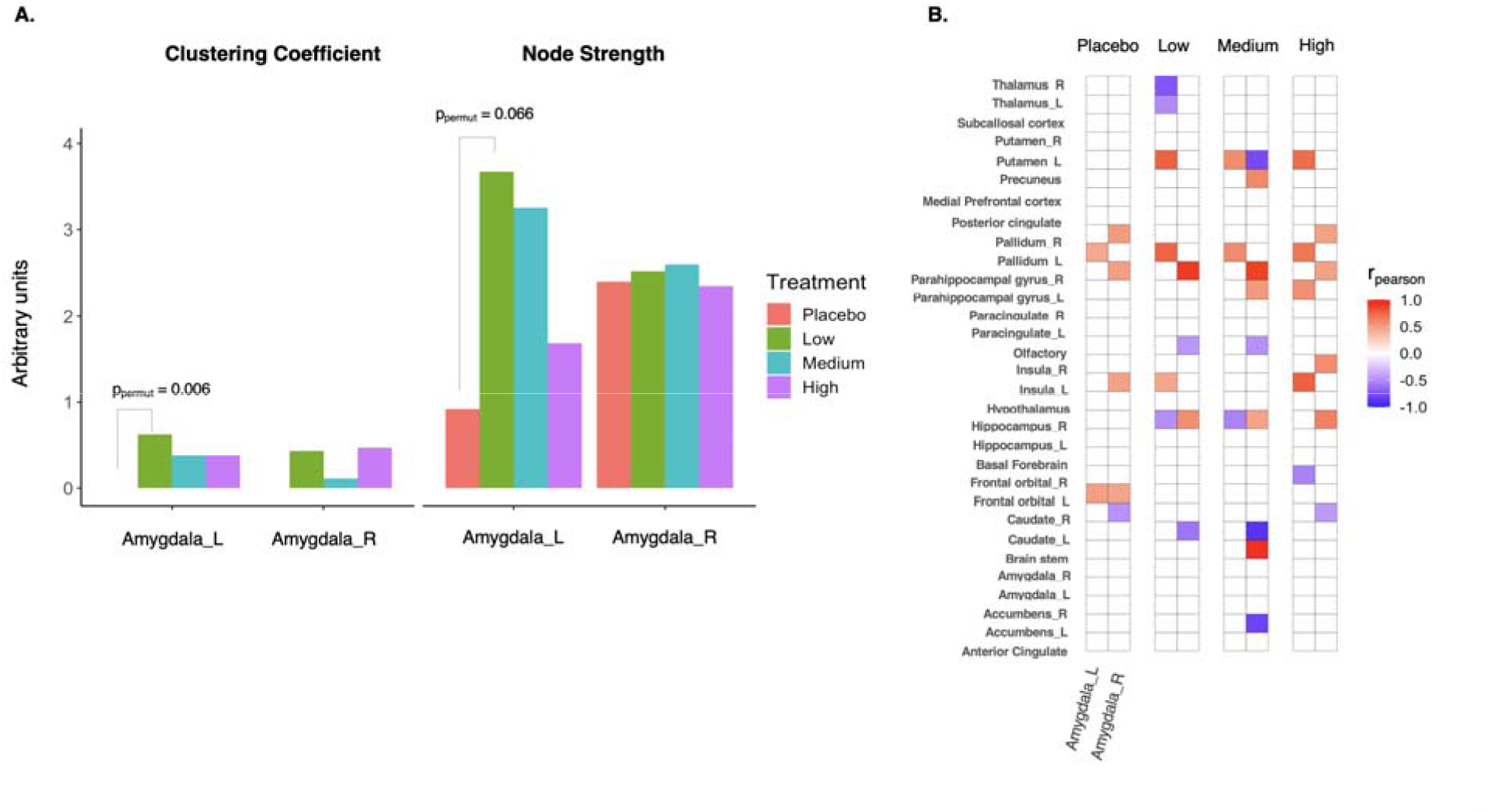
Dose-response effects of intranasal oxytocin on the functional connectivity of the amygdala with the remaining regions of the brain oxytocinergic circuits. We investigated the effects of each dose of intranasal oxytocin, when compared to placebo, on the functional connectivity of the amygdala with the remaining regions-of-interest (ROI) of our brain oxytocinergic network. Instead of conducting statistical analyses on each ROI-to-ROI connections of our network, we summarized the properties of the amygdala’s connections using two graph-theory modelling metrics, node strength and clustering coefficient. Then, we compared node strength and clustering coefficient between each dose and placebo, using permutation testing (10000 permutations). Statistical significance was set to *p* < 0.05 (two-tailed). When a significant effect from placebo was found, we then compared each pair of doses directly. In panel A, we plot the clustering coefficient and node strength of the right and left amygdala for each treatment level. In panel B, we show for illustrative purposes a heatmap of the significant ROI-to-ROI rCBF correlations (p<0.05) involving the right and left amygdala (non-significant correlations were kept white) to help us to understand what changes in functional connectivity are more likely to be driving the changes in clustering coefficient and node strength we present for the low dose in panel A. Colours represent the magnitude of the correlation coefficient; R – Right; L – Left.

To help us interpret the changes in the clustering coefficient and node strength we found for the left amygdala after the low dose compared to placebo, we show for illustrative purposes in Figure 3B all significant correlations between the rCBF in the right and left amygdala and all remaining areas of our oxytocinergic network, for each treatment level separately. This figure indicates that the changes in node strength and clustering coefficient we found in the left amygdala after the low dose compared to placebo are likely to be driven by the following functional connectivity changes. First, low dose oxytocin increased the functional connectivity between the left amygdala and left pallidum (this connection passed our significance threshold in the placebo session and was increased by low dose oxytocin). At the same time, the low dose oxytocin strengthened several connections that did not reach significance in the placebo session, such as positive correlations between the left amygdala and the left insula and left putamen, and negative correlations with the thalamus bilaterally and right hippocampus.

#### Amygdala (subdivisions)

When we repeated the same analysis using ROIs of the subdivisions of the amygdala instead of one ROI for the whole amygdala, we could not detect any significant effect of treatment on node strength or clustering coefficient of any of the right or left subdivisions ROIs we examined (smallest p_permuted_ = 0.193) (Supplementary Figure 4).

#### Exploratory analysis of functional connectivity changes in the whole oxytocinergic network

We also compared the similarity of the connectivity matrices from each dose and placebo by calculating Pearson’s cross-correlation coefficients. We found that our matrices of the placebo and low dose conditions were not significantly correlated with each other (r = 0.197, p=0.108), while we found significant correlations between placebo and the medium dose (r = 0.247, p=0.032), and between placebo and high dose (r = 0.513, p<0.0001). Numerically, the magnitude of these cross-correlations increased with increasing doses (low < medium < high) (Figure 4). The cross-correlation coefficients for the low and medium doses were significantly lower than the cross-correlation coefficient for the high dose (low vs high: Z = −2.124, p = 0.017; medium vs high: Z = −1.874, p = 0.030). Direct comparisons of the cross-correlation coefficients between the low and medium doses yielded no significant differences (Z = −0.308, p = 0.379). Altogether, the findings from these cross-correlation similarity analyses suggested that the effects of intranasal oxytocin on the functional connectivity of the brain’s oxytocinergic circuit were maximal for the low dose and returned to placebo levels as dose increased.

**Fig. 4.**
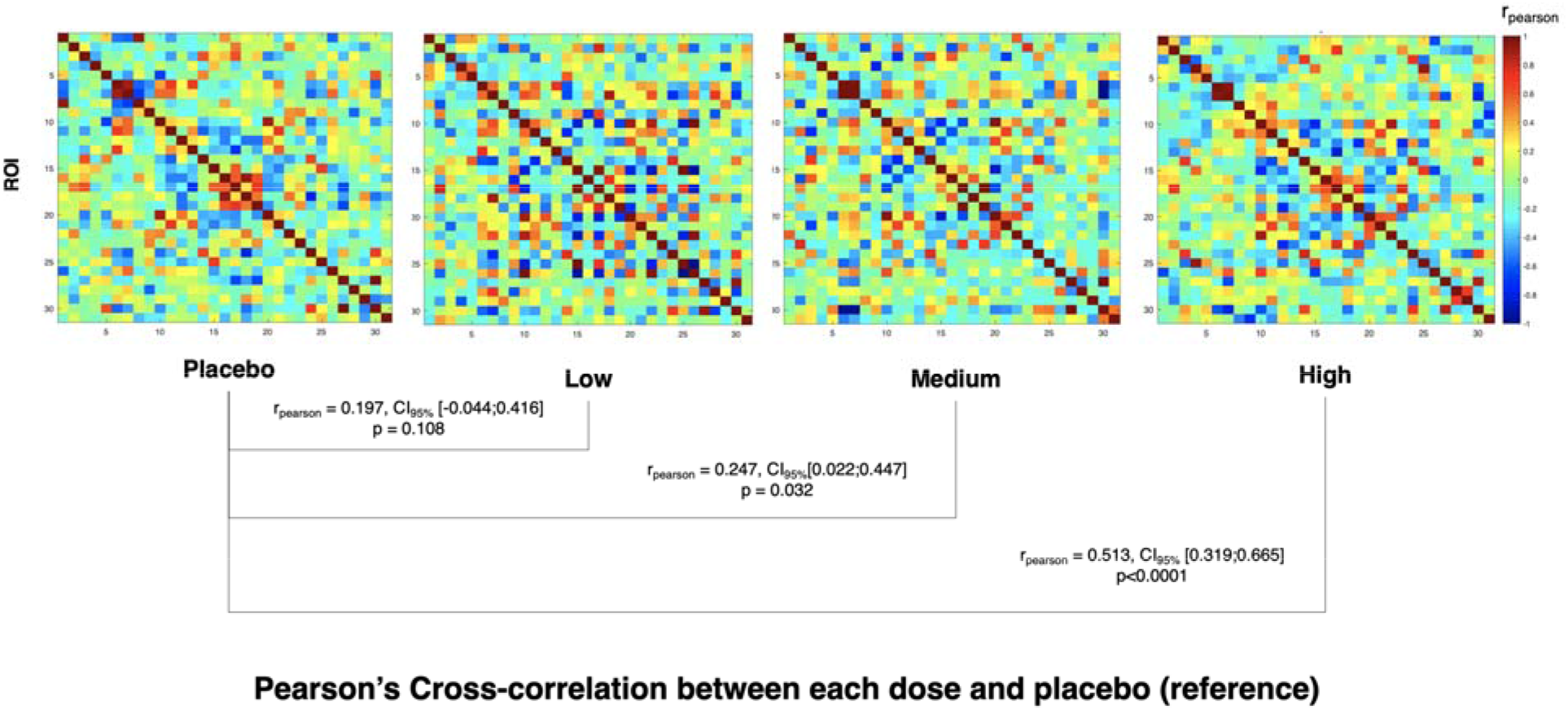
Dose-response effects of intranasal oxytocin on the functional connectivity of the oxytocinergic circuits in the human brain. We investigated the effects of our different doses of intranasal oxytocin on the functional connectivity of an oxytocinergic network encompassing 31 regions-of-interest (ROI) suggested to be part of the main oxytocinergic circuits in the human brain. We used group-based regional cerebral blood flow (rCBF) covariance as a proxy for functional connectivity between ROIs. For each of our four treatment conditions, we created rCBF-covariance matrices reflecting the correlations of rCBF between each pair of ROIs of our oxytocinergic network across subjects. We show in this figure these four symmetric 31 x 31 ROIs matrices. We then assessed treatment related-effects using Pearson’s cross-correlations between the lower triangles of each of our three doses covariance matrices and the one from placebo (reference) as a measure of between-matrices similarity. Please note that these cross-correlations were calculated using only the significant correlations (p < 0.05) present in the two matrices of each pair (i.e. elements of the matrices that overlap after thresholding) – we did not include all other non-significant correlations to reduce noise from potential spurious correlations. In simple terms, a significant high cross-correlation is indicative of high similarity between the treatment and placebo matrices. This would be compatible with absence of significant treatment effects. Decreases in cross-correlations coefficients across doses indicate decreases in similarity with the placebo’s matrix and therefore increases in the magnitude of treatment effects for a certain dose. For each of these three cross-correlations we report the Pearson’s correlation coefficient *(r_pearson_)* and the correspondent 95% confidence interval and *p*-value. Statistical significance was set to *p* < 0.05.

Repeating the same analyses substituting the whole amygdala ROIs for the amygdala subdivisions ROIs in the oxytocinergic network yielded similar findings (see Supplementary Figure 5 for further details).

### Relationship between intranasal oxytocin-induced changes in rCBF and the expression of OXTR mRNA in the post-mortem human brain

To investigate our second main question, we examined whether the expression of the mRNA of the *OXTR* in the post-mortem human brain could predict the magnitude of intranasal oxytocin-induced changes in rCBF (ΔCBF) across regions of the whole brain for each of the three doses we tested here. In Figure 5, we provide brain maps depicting the regional distribution of the *OXTR* mRNA in the *post-mortem* human brain and of the ΔCBF induced by each dose of intranasal oxytocin (Figure 5). We found significant negative correlations between the mRNA expression of the *OXTR* gene and the rCBF changes induced by the low (spearman rho = −0.53, p_spin_<0.001) and medium (spearman rho = −0.43, p_spin_ = 0.002) doses. These correlations were not significant for the high dose (spearman rho = −0.17, p_spin_ = 0.259). We noticed a pattern of nominally decreasing strength of these correlations with increasing dose (Low > Medium > High) (Figure 6A and B). However, direct comparisons of these mRNA-ΔCBF correlations between the three active doses groups yielded no significant differences (Z=-0.507, p=0.306). For both the low and medium doses, the *OXTR* gene was among the strongest negatively correlated of all genes (Figure 6C).

**Fig. 5.**
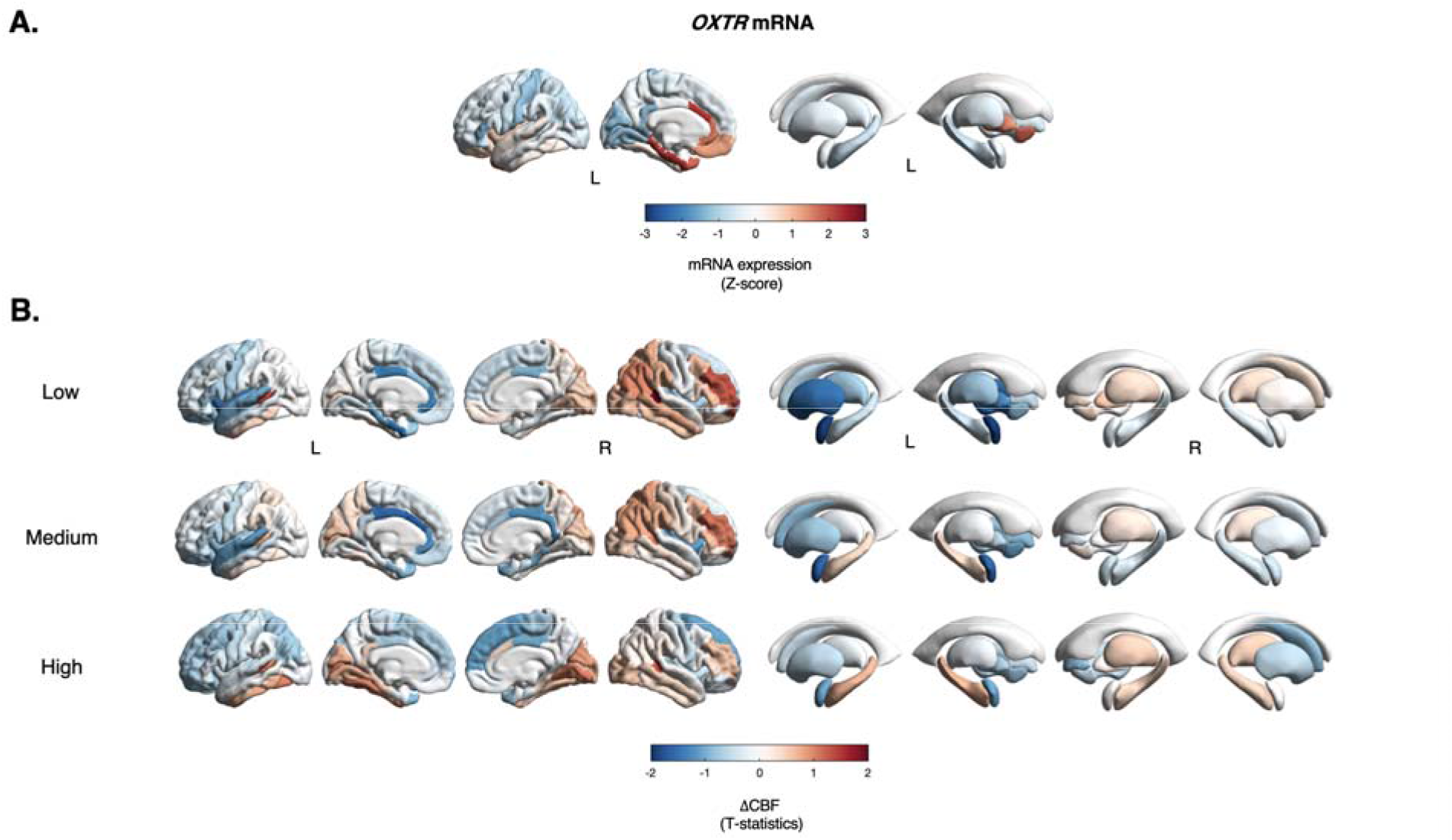
Regional distribution of *OXTR* mRNA in the *post-mortem* human brain and changes in perfusion induced by each dose of intranasal oxytocin. In panel A, we present a brain map showing the spatial distribution of the mRNA of the oxytocin receptor (*OXTR*) in the post-mortem human brain (microarray data from the *Allen Brain Atlas* (ABA)) mapped to the Desikan-Killiani atlas (left hemisphere only). Colours depict z-score of mRNA expression. In panel B, we present the changes in CBF induced by each dose of intranasal oxytocin as compared to placebo, in each parcel of the DK atlas. Colours depict average T-statistics of all voxels within each region (Dose > Placebo).

**Fig. 6.**
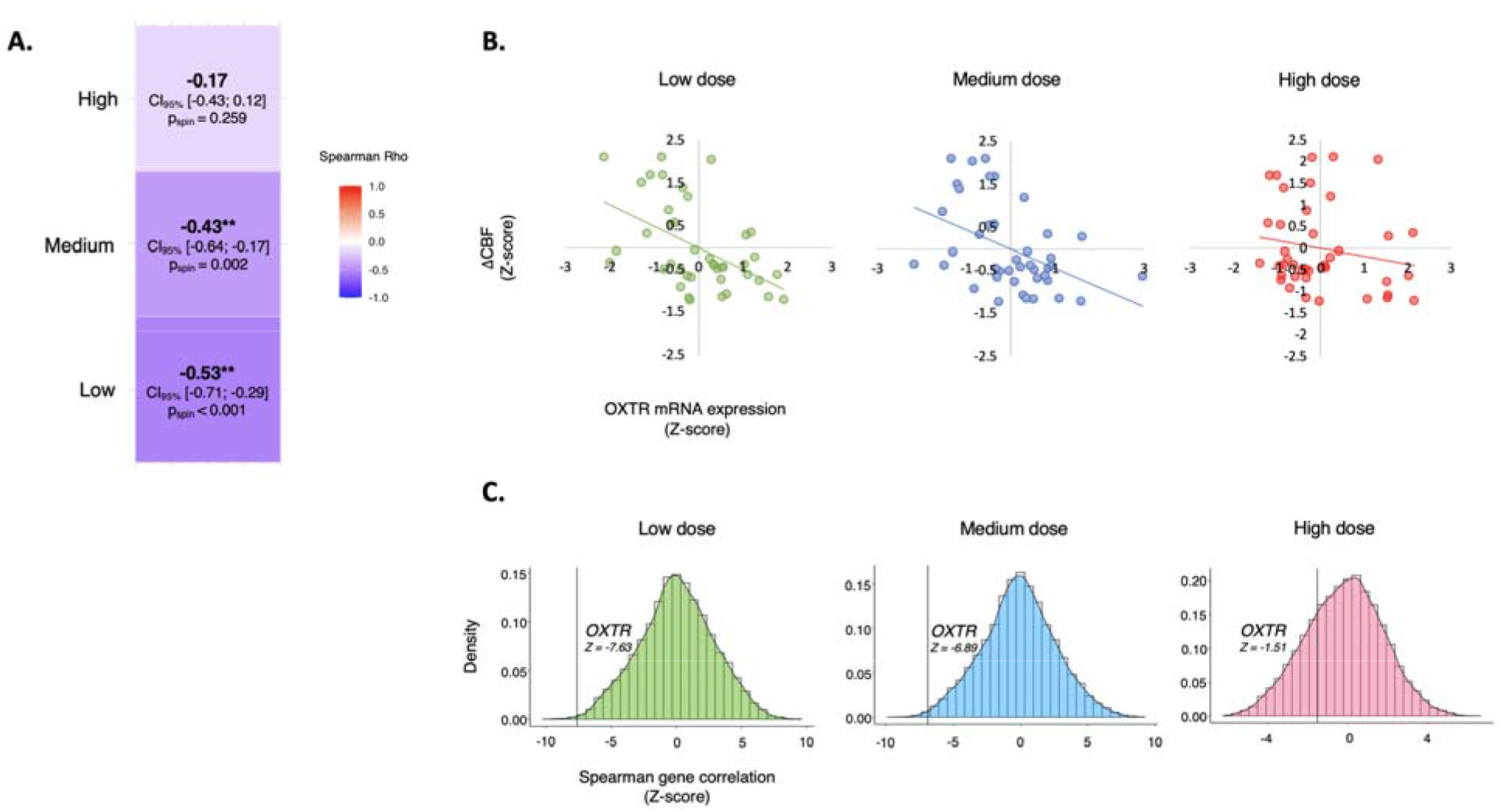
Relationship between intranasal oxytocin-induced changes in regional cerebral blood flow (rCBF) and the mRNA expression of the oxytocin receptor in the post-mortem human brain. In panel A, we present a heatmap showing non-parametric Spearman correlations between each intranasal oxytocin dose-induced changes in rCBF and the levels of mRNA of the oxytocin receptor (OXTR) in the *post-mortem* human brain (microarray data from the *Allen Brain Atlas* (ABA)). Colours depict the magnitude of the Spearman’s correlation coefficient. Statistically significant correlations (*p* <0.01) are highlighted using two asterisks **. In panel B, we present scatter plots for each of these correlations (*X-axis* – *Z*-scores of OXTR’s mRNA expression levels; *Y-axis* – *Z*-scores of each dose-associated changes in rCBF from placebo (ΔCBF); each dot represents one of the 41 regions-of-interest (ROI) of the DK atlas we used in this analysis to resample both the mRNA and ΔCBF data in the same space). The line presents the result of a linear regression fit line. In panel C, we show density plots depicting distributions of the spearman correlations between all ABA genes that passed our pre-processing criteria (15,633 genes) and the changes in CBF associated with each dose. The black line marks the z-score of the correlation for the *OXTR* gene in the distribution.

### Dose-response effects of intranasal oxytocin on cerebrovascular reactivity

Finally, to investigate our third question, we examined dose-response effects of intranasal oxytocin on cerebrovascular reactivity using the BOLD fMRI data of our breath-hold task. We started by conducting some control analyses to discard potential dose-response effects on respiratory belt signals, which index aspects of the respiratory dynamics of our participants while performing the task, as potential confounders. Here, we focused on the readings of our respiratory belt right at the start of each hold block as an estimate of the amplitude of the last forced exhalation and on the respiratory frequency during the paced breathing blocks. Indeed, we did not find a treatment effect on the readings of our respiratory belt right at the start of each hold block (F(2.921, 64.271) = 0.407, p = 0.743) (Supplementary Figure 6) or on the respiratory frequency during the paced breathing blocks (F(2.349, 51.672) = 0.255, p = 0.810) (Supplementary Figure 7). Therefore, potential treatment effects on CVR cannot be accounted by treatment effects on the respiratory dynamics. Then, we examined whether our task elicited the intended increases in the BOLD signal across the whole brain that is characteristic of the cerebrovascular reactivity contrasts. As expected, we found that hold blocks, as compared to paced breathing, produced widespread increases in the BOLD signal across the whole-brain (Supplementary Figure 8). We could not find any cluster where the BOLD signal during hold was lower than during paced breathing. Finally, we investigated dose-response effects on cerebrovascular reactivity at the whole-brain level. We did not find any cluster showing a significant treatment effect on CVR.

## Discussion

Our study contributes three key novel insights regarding the pharmacodynamics of intranasal oxytocin in the human brain. First, using a novel device that maximizes deposition in putative areas of nose-to-brain transport and a wide range of doses, we demonstrate that intranasal oxytocin-induced changes in local rCBF in the amygdala at rest, and in the covariance between rCBF in the amygdala and other key hubs of the brain oxytocin system, follow a dose-response curve with maximal effects for lower doses. These effects were driven by regional effects in the centromedial amygdala and the amygdalostriatal transition area, where we found a similar dose-response pattern in each subregion individually. However, we also demonstrate that a specific amygdala sub-region (superficial amygdala) responds to oxytocin with a different dose-response profile, which highlights the need to qualify the selection of dose by targeted subdivision for maximal pharmacologic effects. Second, we demonstrate that intranasal oxytocin-induced changes in brain’s physiology are linked with inter-regional differences in the density of mRNA levels of the *OXTR* gene across the human brain. Third, we demonstrate that intranasal oxytocin does not disrupt cerebrovascular reactivity in doses between 9-36 IU, supporting the validity of using hemodynamic MRI markers to probe its effects on the human brain in the absence of major vascular confounds. We discuss each of these three main findings below.

Our first key finding was the observation that rCBF changes induced by intranasal oxytocin in the amygdala show a dose-response pattern that varies by subdivision. For most subregions smaller doses produce the largest decreases in rCBF and exceeding this optimal dose-range (9-18 IU) resulted in smaller or null effects. We demonstrate this dose-response profile in a hypothesis-driven analysis of local changes in rCBF in two subdivisions of the left amygdala, the centromedial amygdala and the amygdalostriatal transition area, and in measures of functional connectivity between the left amygdala and other key brain areas of the central oxytocinergic circuit (such as the basal ganglia or the insula). Further, we show that this pattern extends beyond the amygdala and reflects in changes in the inter-regional functional connectivity of a network of brain areas that includes key nodes of the central oxytocinergic circuits. However, we also found that decreases in rCBF in the superficial amygdala can only be produced by higher doses (36 IU), but not lower doses. The independent modulation of physiology (rCBF) in amygdala’s subdivisions by a single dose of intranasal oxytocin, compared to placebo, that we observed in this study corroborates previous studies using BOLD-fMRI^50, 66, 67^. Here, we extend these previous findings by using a physiological quantitative neuroimaging biomarker that can be more closely linked to changes in neural activity and by characterizing the dose-response profile of each subdivision individually. By doing so, we highlight the dose-response profile of perfusion changes following intranasal oxytocin in different amygdala subdivisions is complex and needs to be qualified within each subdivision.

The suppression of amygdala’s activity constitutes one of the most robust findings in animal studies and intranasal oxytocin studies in men^44–47^. The decrease in resting rCBF in the left amygdala after intranasal oxytocin in the current study is consistent with our previous within-subject^22^ report. The precise neural mechanisms underlying the decrease in amygdala rCBF induced by intranasal oxytocin remain unclear. Nevertheless, our findings dovetail with previous observations that oxytocin can inhibit neurons in the centromedial amygdala – probably through its excitatory effects on γ-aminobutyric acid inhibitory (GABAergic) local projections that originate in the lateral and capsular parts of the central nucleus of the amygdala^49, 66^.

Why might the amygdala subdivisions show distinct dose-response profiles, i.e. respond differently to the same nominal doses of intranasal oxytocin? It is possible that the differences in the dose-response profile between the centromedial amygdala subdivision and the amygdalostriatal transition area, on one hand, and the superficial subdivision, on the other hand, that we report here could reflect differences in the expression of oxytocin targets across these subdivisions. To gain insight on this hypothesis, we used transcriptomic data from the Allen Brain Atlas to quantify mRNA expression of the *OXTR* and *V1aR* genes in the centromedial, laterobasal and superficial amygdala subdivisions (expression data for the amygdalostriatal transition area was not available). While the number of donors was too small for a meaningful statistical analysis, we noted that for 4/6 donors mRNA expression of these two genes is consistently lower in the left superficial amygdala than in the left centromedial and laterobasal amygdala subdivisions (Supplementary Figure 9). If the expression of oxytocin targets is indeed lower in the superficial amygdala than in the other subdivisions, this could justify why only a larger dose decreases rCBF in this specific subdivision.

The exact mechanisms explaining the dose-response profile of a given brain region remain unclear. Apart from differences across regions in the level of expression of oxytocin targets, we believe that at least three other mechanisms can play a critical role. All three mechanisms reflect differences in the sensitivity of the cellular signalling processes as a function of the concentration of oxytocin in the extracellular fluid. First, a higher dose of oxytocin and, therefore, the resulting increased oxytocin concentration in the extracellular fluid, may: (i) recruit differentially G_q_/G_i_ pathways when binding the OXTR^24, 25^. A higher concentration of oxytocin induces the recruitment of the inhibitory G_i_ pathway which counteracts the effects on the stimulatory G_q_ pathway, resulting in decreased, null, or even opposite oxytocin effects; (ii) induce the activation of vasopressin receptors and counteract OXTR-binding related effects. This mechanism has been supported by one rodent study in relation to the effects of oxytocin on the amygdala^27^; (iii) result in fast oxytocin receptors internalization^24^, at least in some brain regions. *In-vitro* studies have shown that this process can happen as quickly as 5-15 mins after exposure to oxytocin^68–70^. However, testing these hypotheses directly is virtually impossible in humans for now.

Our findings highlight the need to carefully consider dose in our efforts to engage central oxytocinergic circuits in the living human brain. In the absence of dose-response evidence for specific outcomes, our study corroborates previous reports^30–35^ in identifying lower doses as the most likely effective starting-point. However, given the complexity of the oxytocin signalling machinery and the inter-region differences in the expression of oxytocin’s targets, it is plausible that different brain regions might respond to intranasal oxytocin with different dose-response profiles. We show this for a specific subdivision of the amygdala; future and larger studies should focus on expanding our understanding of the dose-response profile of other key regions of the oxytocinergic network. Ultimately, this information could optimize the engagement of target brain regions for specific applications.

Our second key finding was the observation that rCBF changes for the low and medium doses can be predicted by the distribution of the *OXTR* mRNA expression in the post-mortem human brain. This provides indirect but supportive evidence for a link between the brain’s functional changes that follow the administration of intranasal oxytocin in the living human brain and OXTR engagement. While doubts persist about the exact mechanisms through which oxytocin may reach and induce functional effects in the brain when administered intranasally^71^, our findings provide an extra level of evidence strengthening our confidence on the utility of intranasal administrations of oxytocin as a valid method to target the oxytocin system in the human brain. However, our indirect findings should not detract future studies combining the administration of intranasal oxytocin with brain-penetrant antagonists, once these antagonists become widely available in the future, to validate our findings further. Certainly, it would have been interesting to examine whether the distribution V1aR mRNA could also explain some of the variance in the changes in brain’s physiology that follow intranasal oxytocin and to test whether this predictive ability might increase for higher doses of intranasal oxytocin (which more likely engage this AVP receptor). However, for now, testing this hypothesis with the ABA data will be impossible given that the mRNA of V1aR could not be reliably measured above background.

Our third key finding was the lack of effects of intranasal oxytocin on cerebrovascular reactivity. Even though oxytocin is a vasoactive neuropeptide^61^ and unspecific effects of oxytocin on cerebrovascular reactivity would undermine the validity of using neuroimaging measures as biomarkers of the brain responses to intranasal oxytocin^60^, this important methodological question has been left unaddressed over the years. Our study shows that intranasal oxytocin does not disrupt cerebrovascular reactivity across a range of doses. It therefore strengthens our confidence on the validity of using indirect physiological measures of brain function, such as rCBF or BOLD changes, to probe the effects of intranasal oxytocin on the living human brain, in the absence of major unspecific vascular confounders.

Our study faces certain limitations. First, our findings cannot be readily extrapolated to women, given the known sexual dimorphism of the oxytocin system^72–74^. Second, in this study we administered intranasal oxytocin using the PARIS SINUS nebulizer, which increases deposition in the regions of the upper nose putatively involved in the nose-to-brain transport of oxytocin^75, 76^. While this does not detract from the dose-response profile we present here, it may make direct comparisons with nominal doses delivered with other devices for nasal delivery, including standard sprays which may be less efficient in oxytocin delivery^77^, challenging. Third, in this study we sampled rCBF at 14-32 mins post-dosing based on our previous work showing that changes in amygdala rCBF after oxytocin, despite route of administration, emerge already at 15-32 minutes post-dosing^22^. While our choice of time-interval served the aims of the current study, future studies should perform a comprehensive characterization of the spatiotemporal profile of the changes in rCBF that follow the different doses we administered here over a longer period. Fourth, given that our sample consisted of healthy participants, our findings might not directly extrapolate to clinical groups that may present disease-related changes in oxytocin signalling (i.e. decreases or increases in OXTR expression)^78–81^ that shift intranasal oxytocin’s response curve. Fifth, we used the distribution of the mRNA levels of the *OXTR* gene in the post-mortem human brain as a proxy for its levels in the living human brain. Since post-transcriptional events may alter the relationship between gene expression and protein synthesis^82^, it would be important to ascertain the validity of using mRNA levels as proxies for the OXTR proteins in the living human brain (this is virtually impossible to verify until a PET ligand for OXTR becomes available). Finally, while we have conducted our analysis of the effects of the different doses of intranasal oxytocin on CVR at the whole-brain level, our coverage of areas showing BOLD signal dropout due to MRI susceptibility effects, such as the subcallosal and orbitofrontal cortices or the ventral striatum, during the breath hold task, was relatively poor. Therefore, we cannot discard with confidence potential effects on CVR in these specific brain areas – which are often reported to be modulated by intranasal oxytocin^83, 84^.

In conclusion, our data highlight the need to carefully consider dose in our efforts to engage central oxytocinergic circuits in the human brain by using intranasal oxytocin and suggests that a one-size-fits-all approach might not capture differences in dose-response between target regions. Furthermore, it strengthens our confidence in the validity of using intranasal oxytocin to target the brain’s oxytocin system and indirect MRI-based neuroimaging measures to probe its effects on brain’s function in the absence of major vascular confounders.

## Methods

### Participants

We recruited 24 healthy male adult volunteers (mean age 23.8 years, SD = 3.94, range 20-34 years). We screened participants for psychiatric conditions using the MINI International Neuropsychiatric interview^85^. Participants were not taking any prescribed drugs, did not have a history of drug abuse and tested negative on a urine panel screening test for recreational drugs, consumed <28 units of alcohol per week and <5 cigarettes per day. We instructed participants to abstain from alcohol and heavy exercise for 24 hours and from any beverage other than water or food for at least 2 hours before scanning. Participants gave written informed consent. King’s College London Research Ethics Committee (HR-17/18-6908) approved the study. We determined sample size based on our two previous studies demonstrating that N=16 per group was sufficient to quantify intranasal oxytocin-induced changes in rCBF in between-^39^ and within-subject^22^ designs.

### Study design

We employed a double-blind, placebo-controlled, crossover design. Participants visited our centre for 1 screening session and 4 experimental sessions spaced 4.3 days apart on average (SD = 5.5, range: 2-16 days). During the screening visit, we confirmed participants’ eligibility, obtained informed consent, collected sociodemographic data and measured weight and height. Participants also completed a short battery of self-report questionnaires (which are not reported here). Participants were trained in a mock-scanner during the screening visit to habituate to the scanner environment and minimize its potential distressing impact. Participants were also trained on the correct usage of PARI SINUS nebulizer, the device that they would use to self-administer oxytocin or placebo in the experimental visits. Participants were randomly allocated to a treatment order using a Latin square design.

### Intranasal oxytocin administration

Participants self-administered one of three nominal doses of oxytocin (Syntocinon; 40IU/ml; Novartis, Basel, Switzerland). We have consistently shown in two separate studies in healthy men that intranasal oxytocin (40 IU) modulates rCBF in the amygdala^22, 39^. In our last within-subject study investigating the effects of intranasal (40 IU administered using either a standard spray or the PARI SINUS nebuliser) and intravenous (10 IU) administrations of oxytocin on rCBF in healthy men, we showed that suppression of amygdala rCBF is an early post-dosing effect (15-32 mins) that emerges irrespective of route of administration^22^. Hence, in the current study, we selected the same time interval, a range of doses of intranasal oxytocin (9, 18 and 36 IU) or placebo. Placebo contained the same excipients as Syntocinon except for oxytocin. Immediately before each experimental session started, a researcher not involved in data collection loaded the SINUS nebulizer with 2 ml of a solution (1 ml of which was self-administered) containing oxytocin in the following concentrations 40 IU/ml, 20 IU/ml and 10 IU/ml or placebo (achieved by a simple 2x or 4x dilution with placebo).

Participants then self-administered each dose of oxytocin (Syntocinon) or placebo, by operating the SINUS nebulizer for 3 minutes in each nostril (6 min in total), based on a rate of administration of 0.15-0.17 ml per minute. In pilot work using nebulization on a filter, we estimated the actual nominally delivered dose for our protocol to be 9.0IU (CI 95% 8.67 – 9.40) for the low dose, 18.1IU (CI 95% 17.34 – 18.79) for the medium dose and 36.1IU (CI 95% 34.69 – 37.58) for the high dose. The correct application of the device was validated by confirming gravimetrically the administered volume. Participants were instructed to breathe using only their mouth and to keep a constant breathing rate with their soft palate closed, to minimize delivery to the lungs. The *PARI SINUS* nebuliser (PARI GmbH, Starnberg, Germany) is designed to deliver aerosolised drugs to the sinus cavities by ventilating the sinuses via pressure fluctuations. Droplet diameter is roughly one tenth of a nasal spray and its mass is only a thousandth. As a result, the SINUS nebuliser can increase delivery to sinuses, upper nose and olfactory regions^75^. One study has shown up to 9.0% (±1.9%) of the total administered dose with the SINUS nebuliser to be delivered to the olfactory region, 15.7% (±2.4%) to the upper nose; for standard nasal sprays, less than 4.6% reached the olfactory region^76^.

Participants guessed the treatment condition correctly on 24 out of the total 96 visits (25%), which was not significantly different from chance (Chi-squared test: χ^2^ (9) = 15.83, p = 0.070) (Supplementary Table 3).

### Procedure

All participants were tested at approximately the same time in the afternoon (3-5 pm) for all oxytocin and placebo treatments, to minimise potential circadian variability in brain activity^86^ or oxytocin levels^87^. Each experimental session began with a quick assessment of vitals (blood pressure and heart rate) and the collection of two 4 ml blood samples for plasma isolation (data not reported here). Then we proceeded with the treatment administration protocol that lasted about 6 minutes in total (Fig. 1). Immediately before and after treatment administration, participants completed a set of visual analog scales (VAS) to assess subjective drug effects (alertness, mood and anxiety). After drug administration, participants were guided to an MRI scanner, where we acquired a BOLD-fMRI scan during a breath-hold task (lasting 5 minutes 16 seconds), followed by 3 pulsed continuous ASL scans (each lasting 5 minutes and 22 seconds), a BOLD-fMRI scan during a prosocial reinforcement learning task and a resting-state BOLD-fMRI scan. In this manuscript, we report only the results from the analysis of the breath-hold and ASL data (remaining data to be reported elsewhere). We describe the details of each of these two types of scan below. When the participants left the MRI scanner, we assessed subjective drug effects using the same set of VAS.

### MRI data acquisition

We acquired the MRI data in a MR750 3 Tesla GE Discovery Scanner (General Electric, Waukesha, WI, USA) using a 32-channel receive-only head coil. We recorded physiological data using a respiratory belt (for breathing rate) wrapped around the diaphragm and a pulse oximeter (for heart rate) placed on the index or middle finger of the left hand of our participants.

### Anatomical image acquisition

We acquired a 3D high-spatial-resolution, Magnetisation Prepared Rapid Acquisition (3D MPRAGE) T1-weighted scan. Field of view was 270 x 270 mm, matrix = 256 x 256, 1.2 mm thick slices, TR/TE/TI = 7328/3024/400 ms, flip angle 11°,. The final resolution of the T1-weighted image was 1.1 x 1.1 x 1.2 mm.

### Breath-hold task

The breath hold paradigm provides a non-invasive, quick and reliable way of assessing CVR^65, 88^. As CO_2_ is a vasodilator, the hypercapnia induced by holding one’s breath increases the concentration of CO_2_ in the blood^65^, which induces widespread increases in CBF, increasing the BOLD signal across the brain^65^. This increase in the BOLD signal can be used as a proxy for CVR in the absence of neural activity^65^. The breath hold paradigm has been shown to be comparable to other direct methods for assessment of CVR, such as controlled CO_2_ inspiration ^89^. Furthermore, it has been shown to capture cerebrovascular alterations related to disease states^90^, healthy aging^91^, or unspecific effects of pharmacological compounds^60, 92^. Therefore, we used the breath hold task in this study to investigate whether different doses of intranasal oxytocin might disrupt CVR.

Our acquisitions for the breath hold task started at about 8 (±1) mins post-dosing (the earliest time possible allowing for setting up the participant in the scanner) and lasted 5 mins and 16 seconds. We chose to sample this specific time-interval because unspecific effects of intranasal oxytocin on CVR are more likely to occur when concentrations of oxytocin in circulation are maximal and we have previously shown oxytocin to peak in the plasma in the first 15 mins immediately after intranasal oxytocin administration^22^. Therefore, sampling CVR at this time-interval would maximize our chances of capturing intranasal oxytocin-induced effects on CVR, if they existed. Participants were instructed to follow a simple set of instructions on screen alternating between paced breathing (45 seconds) and breath holding (16 seconds), repeating this cycle four times. The task started and ended with a period of paced breathing. Participants were instructed to commence breath holding at the end of a forced expiration, which has been shown to produce a quicker CVR peak, in addition to removing some of the confounds produced by an end-inspiration approach, such as a biphasic response within the time course signal and marked inter-subject variability in inspiration depth^65^. During the regular breathing portions of the task, participants were given instructions to breath at a controlled rate (breath in for 3 seconds, out for 3 seconds) as this approach produces a larger peak of the BOLD signal and improved signal-to-noise ratio than self-paced breathing^93^. Finally, as the BOLD increase during breath hold has been shown to plateau around 20 seconds^94^, we chose a 16 second hold to be long enough to produce a peak response whilst not being uncomfortable for the participant (and not increasing the likelihood of head movement). Standardised verbal reminders of the instructions were given prior to entering the scanner and immediately before the task started. Participants were advised to not breath more heavily or deeply than they normally would during the paced breathing segments; rather they should just breath regularly but in time with the instructions, which were presented visually during the task. We recorded participants’ chest movement using a respiratory belt sensor and monitored their breathing from the control room to ensure they were following the task correctly.

We acquired these functional scans using T2*-sensitive gradient echo planar imaging optimised for parallel imaging ([TR] = 2000 ms; echo time [TE] = 28 ms; flip angle = 75°; field of view = 240 x 240 mm, matrix = 64 x 64, 3 mm slice thickness with a 0.3 mm slice gap, 41 slices, voxel size = 3.75 x 3.75 x 3 mm).

### Arterial Spin Labelling

We performed three 3D pseudo-continuous Arterial Spin Labelling (3D-pCASL) scans to measure changes in rCBF over 14-32 min post-dosing. We sampled this specific time-interval because we have previously shown that the intranasal administration of OT (40 IU) using either a standard nasal spray or the PARI SINUS nebulizer results in robust rCBF changes 15-32 mins post-dosing^22^. Participants were instructed to lie still and maintain their gaze on a centrally placed fixation cross during scanning.

The 3D-pCASL sequence was acquired with a fast spin echo (FSE) stack of spiral readout. We used the following parameters: 10 spiral arms, 600 points per arm, in-plane resolution = 2.94 x 2.94 mm^2^, slice thickness = 3 mm, 54 slices, label duration (LD) = 3500 ms, post-labelling delay (PLD) = 2025 ms, TE/TR = 11.8 / 7325 ms, number of averages = 2, total time of acquisition = 5 mins and 22 seconds. Using a long PLD of 3500ms allowed us to increase the volume of labelled arterial blood and hence maximize SNR. The readout resolution provided us with a better chance to investigate small brain regions, such as amygdala subdivisions. We applied a background suppression module to null static signal, using four inversion pulses. This consisted of one single selective saturation pulse applied to the imaging volume and an imaging selective inversion pulse prior to labelling followed by three non-selective inversion pulses between the end of the labelling block and the readout window. We set the labelling plane to 2 cm below the imaging volume. The imaging volume was positioned on the inferior surface of the cerebellum for all subjects. We also acquired a 3D proton density (PD) image using identical readout parameters for CBF quantification and to aid co-registration. Computation of CBF was done according to the formula suggested in the recent ASL consensus article^95^.

### MRI data preprocessing

#### Breath-hold task

We pre-processed our functional scans using a standard pipeline which included slice time correction, realignment, co-registration to each individual’s structural scan, normalisation to the Montreal Neurological Institute (MNI) 152 standard template and smoothing using a 6 mm FWHM isotropic kernel. We did not have to exclude any participant because of excessive movement (the mean frame-wise displacement for all scans was < 0.5 mm).

We then proceeded with first-level analysis and used the onsets of the paced breathing and breath holding blocks to construct two regressors for which event timings were convolved with a canonical haemodynamic response function and its temporal and shape form derivatives. Since the block design of this task relies on accumulation of CO_2_ over time in the blood, we delayed the onset of the blocks regressors by 9 seconds (hence we used event durations: Paced = 36 seconds; Hold = 7 seconds). This modelling approach was previously shown by *Murphy et al. (2011)* to increase sensitivity as compared to a simple block design without the onset delay of the block regressors^96^. We also included the six head motion realignment parameters to model the residual effects of head motion as covariates of no interest. Data were high-pass filtered at 128s to remove low-frequency drifts. This first-level analysis resulted in the creation of CVR contrast images quantifying BOLD-changes associated with breath hold as compared to paced breathing (Hold vs Paced) for each participant/session. These CVR contrasts were then entered into second-level group statistical analysis to examine task and treatment effects (as described below in the Statistical analysis section). Preprocessing, first and second level analyses were performed using Statistical Parametric Mapping (SPM) 12 (http://www.fil.ion.ucl.ac.uk/spm/software/spm12/).

### Arterial Spin Labelling

A multi-step approach was performed for the spatial normalization of the CBF maps to MNI space: (1) co-registration of the PD image from each scan to the participant’s T1-image after resetting the origin of both images to the anterior commissure. The transformation matrix of this co-registration step was then used to register the CBF map from the corresponding scan with the T1-image; (2) unified segmentation of the T1 image; (3) elimination of extra-cerebral signal from the CBF maps, by multiplication of the “brain only” binary mask obtained in step[2], with each co-registered CBF map; (4) normalization of the subject’s T1 image and the skull-stripped CBF maps to the MNI152 space using the normalisation parameters obtained in step[2]. Finally, we spatially smoothed each normalized CBF map using an 8-mm Gaussian kernel. All of these steps were implemented using the ASAP (Automatic Software for ASL processing) toolbox (version 2.0)^97^. The resulting smoothed CBF maps were then entered into SPM12 for group-level statistical analysis, as described below.

### Physiological data acquisition and processing

We continuously monitored respiratory movements during all scans using a respiratory belt sensor. Respiratory movement signals were first manually checked for artifacts and low-pass filtered with a fourth-order, Butterworth zero-phase filter (cut-off frequency = 2 Hz). Then, we used a technique involving cross-correlation of the filtered respiratory signal with sinusoidal signals of different frequencies to estimate the time-varying frequency of the respiration as suggested by *Chuen et al. (2016)*^98^. The script we used to implement this analysis can be found at https://github.com/finn42/RespirationTracking.

### Statistical analyses

#### Subjective drug effects

We quantified alertness, mood and anxiety using a set of 16 VAS tapping into these three latent constructs. We confirmed the inner structure of our set of VAS using principal component analysis applied to the baseline (before oxytocin administration) data from each of our four experimental sessions. We derived scores of alertness, mood and anxiety by simply averaging the scores of the respective VAS for each of these three subscales (the items used to calculate each of these subscales can be found in Supplementary Table 4). We investigated the effects of treatment, time and treatment x time on alertness, mood and anxiety scores in a full factorial linear mixed model, including treatment, time-interval and treatment x time-interval as fixed effects, participants as a random effect. This analysis was implemented in SPSS 24. When a significant effect was found, we followed with post-hoc tests, applying Sidak’s correction for multiple comparisons.

### Global CBF Measures

We extracted median global CBF values within an explicit binary mask for grey-matter (derived from a standard T1-based probabilistic map of grey matter distribution by thresholding all voxels with a probability > 0.20) using the *fslstats* command implemented in the FSL software suite. We tested for the effects of treatment, time-interval and treatment x time-interval on global CBF signal in a repeated measures analysis of variance implemented in SPSS 24 (http://www-01.ibm.com/software/uk/analytics/spss/), using the Greenhouse-Geisser correction against violations of sphericity. When a significant effect was found, we followed with post-hoc tests, applying Sidak’s correction for multiple comparisons.

### Dose-response effects of intranasal oxytocin on rCBF at rest

We tested the effects of treatment, time-interval and treatment x time-interval on median rCBF values extracted from unsmoothed CBF maps using *fslstats* in 10 anatomical amygdalar regions-of-interest (ROIs), including: i) the left and right whole amygdala, as primary outcomes; and ii) its respective centromedial, laterobasal, superficial and amygdalostriatal transition area subdivisions, as secondary outcomes. We decided to consider right and left homologous amygdala structures separately as we have previously consistently described left lateralization of the effects of intranasal oxytocin on rCBF in men in two separate studies^22, 39^. Whole and non-overlapping ROIs for the amygdala subdivisions were created from cytoarchitectonically defined probabilistic maps available in the Anatomy toolbox (Institute of Neuroscience and Medicine, Julich, Germany) by thresholding each probabilistic map to include only voxels with a probability of belonging to a certain subdivision higher than 0.80 and binarizing them at the end. We confirmed that these ROIs did not contain voxels belonging to more than one subregion. The final number of unique voxels in each ROI can be found in Supplementary Table 5. We used a full factorial linear mixed model, including treatment, time-interval and treatment x time-interval as fixed effects, participants as a random effect and global grey-matter CBF as a nuisance variable. These analyses were implemented in SPSS24. When a significant effect was found, we followed with post-hoc tests, applying Sidak’s correction for multiple comparisons. Additionally, we also contained the false-discovery rate for the number of ROIs tested at α=0.05 using the Benjamini-Hochberg procedure^99^.

We also conducted a whole-brain exploratory investigation of treatment, time-interval and treatment x time-interval effects on rCBF, using global grey-matter CBF as a covariate, which would allow us to explore potential effects beyond the amygdala ROIs. We used cluster-level inference at α = 0.05 using family-wise error (FWE) correction for multiple comparisons and a cluster-forming threshold of p=0.005 (uncorrected). These statistical thresholds were determined *a priori* based on our own work investigating the effects of intranasal oxytocin on rCBF in humans^22, 39^ and are standardly applied in ASL studies measuring rCBF^100–105^. We also tested a cluster-forming threshold of p=0.001 (uncorrected) to be in line with the current recommendations for the statistical analysis of BOLD fMRI^106^, but this did not change the results.

### Dose-response effects of intranasal oxytocin on functional connectivity and network metrics using group-based rCBF covariance

Arterial spin labelling provides rCBF measurements for each voxel of the brain that typically average multiple paired control-label images to increase SNR, hence resulting in relatively poor temporal resolution (at best, we can sample rCBF approx. every 2 minutes because of our segmented readout approach). Therefore, classical functional connectivity analyses that require within-individual correlations of the rCBF time-series across brain regions are not normally possible due to the small number of available time points. A conceptually similar functional connectivity mapping analysis can be conducted by correlating vectors of median rCBF values across participants from each brain region and treatment level (group-based covariance statistics)^52–56^.

We conducted this analysis using the NetPET toolbox (http://www.nitrc.org/projects/netpet_2018/) – a recently developed pipeline for performing covariance statistics and network analysis of brain Positron Emission Tomography (PET) imaging data^107^. We repurposed this tool to conduct our group-based rCBF-covariance analyses^108^. Briefly, first, we averaged our CBF maps across time-intervals for each treatment level to increase SNR^109^. Then, we extracted median rCBF using *fslstats* from a set of 31 anatomical non-overlapping anatomical ROIs selected for their relevance in the brain oxytocinergic circuit (namely, areas showing enrichment of expression of oxytocin pathway related genes^110^ and areas shown to be modulated by pharmacological manipulation of the oxytocin system in non-human animals or humans^111, 112^). The full list of ROIs can be found in Supplementary Table 6 and Figure 3. To eliminate the potential confounding effect of global CBF on these measures, we regressed out the median global CBF from the median rCBF of each of these ROIs in a linear regression model for each treatment level separately and used the standardized residuals in subsequent analyses. We then created interregional adjacency correlation matrices for each treatment level by calculating Pearson’s correlations between the standardized rCBF vectors for each pair of ROIs of our oxytocinergic network. Each vector consisted of the standardized rCBF values across all participants for a given ROI/treatment level. The resulting mathematical objects were four *31* × *31* symmetric weighted matrices, one for each treatment level (where *31* is the number of ROIs used).

Instead of comparing the ROI-to-ROI correlations in our network for each pair of treatment conditions (which would imply a high number of tests and therefore raise concerns about multiple testing), we used graph-based modelling to summarize the core topographic characteristics of these weighted matrices and allow us to assess interactions between regions^113^. We decided to focus our analysis on two graph-metrics: node strength and clustering coefficient. Node strength is the average connectivity of a node and is defined as the sum of all neighbouring link weights^114^. Clustering coefficient is a measure of functional segregation and quantifies the number of connections that exist between the nearest neighbours of a node as a proportion of the maximum number of possible connections^114^. Both node strength and clustering coefficient are calculated in the NetPET toolbox using the Brain Connectivity Toolbox (https://www.nitrc.org/projects/bct/). According to standard practice, the graph metrics were extracted from the rCBF weighted covariance matrices after thresholding to preserve the strongest functional connections (ROI-to-ROI correlations with p<0.05). The full process was then repeated, this time substituting the left and right whole amygdala by its corresponding centromedial, laterobasal, superficial and amygdalostriatal transition area subdivision ROIs. This resulted in symmetric weighted matrices of 37 x 37 ROIs.

We then conducted a set of analysis to investigate treatment-related effects on these weighted matrices and graphs-metrics. First, given our focus on the amygdala, we compared the left and right amygdala’s node strength and clustering coefficient between each dose and placebo (note that because these analyses compare group-based covariance matrices, only two groups can be compared at a time). We assessed statistical significance by permutation testing^115^, using up to 10,000 permutations to generate the null distribution from the data without any a priori hypothesis about the directionality of the effect. Permutations were implemented by swapping the elements between pairs of treatment conditions being compared and using the resulting difference to generate a null-distribution of permuted values. A significant group difference would be found when the true difference between groups falls into the lowest or highest 2.5% of the distribution of the permuted differences between the two conditions. This is equivalent to a two-tailed test. When a significant difference from placebo was identified, we then compared each pair of active doses using the same permutation approach. The same analysis was then repeated for each amygdala subdivision ROI.

Finally, we also conducted an exploratory analysis investigating treatment-related changes on functional connectivity of our oxytocinergic network as a whole. We investigated the similarity between the adjacency matrices of each of our active dose and placebo groups by calculating cross-correlations between each pair of matrices^116^. These cross-correlations were calculated using only significant ROI-to-ROI correlations (p<0.05) in each pair of matrices (i.e., they were calculated based on the elements of the matrices that overlapped after applying the statistical threshold). We did not include non-significant correlations to reduce noise from potentially spurious correlations. In simple terms, to take the two extremes, a significant and high cross-correlation between the matrices of any dose and placebo would be indicative of high similarity, and therefore suggest the absence of substantial treatment effects. In turn, a non-significant and low correlation would indicate low similarity, and therefore suggest the presence of treatment related changes. We then compared these cross-correlations across doses using r-to-Z transformation. All these analyses were then repeated for the matrices including the amygdala subdivisions in the place of the whole amygdala ROIs.

### Relationship between intranasal oxytocin-induced changes in rCBF and the expression of OXTR mRNA

Regional microarray expression data were obtained from six post-mortem brains provided by the ABA (http://human.brain-map.org/) (ages 24–57 years)^117^. We used the *abagen* toolbox (https://github.com/netneurolab/abagen) to process and map the transcriptomic data to 84 parcellated brain regions from the DK atlas^118^. Briefly, genetic probes were reannotated using information provided by *Arnatkeviciute et al., 2019*^119^ instead of the default probe information from the ABA dataset, hence discarding probes that cannot be reliably matched to genes. Following previously published guidelines for probe-to-gene mappings and intensity-based filtering^119^, the reannotated probes were filtered based on their intensity relative to background noise level; probes with intensity less than background in ≥50% of samples were discarded. A single probe with the highest differential stability, ΔS(p), was selected to represent each gene, where differential stability was calculated as^120^:

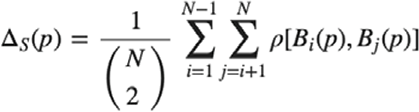

Here, ρ is Spearman’s rank correlation of the expression of a single probe *p* across regions in two donor brains, B_i_ and B_j_, and *N* is the total number of donor brains. This procedure retained 15,633 probes, each representing a unique gene.

Next, tissue samples were assigned to brain regions using their corrected MNI coordinates (https://github.com/chrisfilo/alleninf) by finding the nearest region within a radius of 2 mm. To reduce the potential for misassignment, sample-to-region matching was constrained by hemisphere and cortical/subcortical divisions. If a brain region was not assigned to any sample based on the above procedure, the sample closest to the centroid of that region was selected to ensure that all brain regions were assigned a value. Samples assigned to the same brain region were averaged separately for each donor. Gene expression values were then normalized separately for each donor across regions using a robust sigmoid function and rescaled to the unit interval. We applied this procedure for cortical and subcortical regions separately, as suggested by *Arnatkeviciute et al., 2019*^119^. Scaled expression profiles were finally averaged across donors, resulting in a single matrix with rows corresponding to brain regions and columns corresponding to the retained 15,633 genes. As a further sanity check, we conducted leave-one-donor out sensitivity analyses to generate six expression maps containing gene expression data from all donors, one at a time. The principal components of these six expression maps were highly correlated (average Pearson correlation of 0.993), supporting the idea that our final gene expression maps where we averaged gene expression for each region across the six donors is unlikely to be biased by data from a specific donor. Since the AHBA only includes data for the right hemisphere for two subjects we only considered the 41 regions of left hemisphere regions (34 cortical plus 7 subcortical regions). The V1aR gene did not show levels of expression above background and was excluded from further analyses.

As a proxy for drug effects, we used the whole-brain T-statistical parametric maps comparing each active dose against placebo, which reflect the effect size of the changes in rCBF from placebo associated with each active dose while accounting for global CBF (ΔCBF). These T-statistical maps were calculated by contrasting the averaged CBF maps across time-intervals of each active oxytocin dose level against placebo. We calculated the average T-statistics of these parametric maps in each region of the DK atlas we used to map gene expression.

We then calculated non-parametric Spearman correlations between ΔCBF and the mRNA expression of the *OXTR* gene across the 41 brain regions. Here, we assessed significance using spatial permutation testing (spin test) to account for the inherent spatial autocorrelation of the imaging data, as implemented in previous studies^121–123^. This approach consists in comparing the empirical correlation amongst two spatial maps to a set of null correlations, generated by randomly rotating the spherical projection of one of the two spatial maps before projecting it back on the brain parcel. Importantly, the rotated projection preserves spatial contiguity of the empirical maps, as well as hemispheric symmetry. Past studies using the spin test have focused on comparisons between cortical brain maps. However, subcortical regions were also of interest in this study. Subcortical regions cannot be projected onto the inflated spherical pial surface, so an alternative approach was needed. We incorporated the subcortex into our null models by shuffling the seven subcortical regions with respect to one another, whereas the cortical regions were shuffled using the spin test. We then used r-to-z transformation to compare the correlation coefficients between ΔCBF and mRNA density between doses. Statistical significance was set at p<0.05 (two-tailed).

Finally, we repeated the same procedure for all other AHBA 15,632 genes that passed our pre-processing criteria to assess the specificity of the correlations between ΔCBF and mRNA density we found for the *OXTR* gene. We ranked all genes according to their correlation with ΔCBF associated with each dose of intranasal oxytocin and checked where in the distribution of these correlations the *OXTR* gene lies.

### Breath-hold: dose-response effects of intranasal oxytocin on cerebrovascular reactivity

Changes in the respiratory dynamics induced by intranasal oxytocin could produce differences in the amounts of circulating CO_2_ in the blood during both the hold and paced breathing blocks of our task and confound our CVR assessments^96^. Therefore, before performing any analysis on our CVR contrasts, we conducted some sanity checks using the respiratory data we acquired during the task to dismiss such confounders. First, we examined treatment effects on the respiratory belt readings right at the start of each hold block, which would give us an idea about whether intranasal oxytocin could have changed the amplitude of the last exhalation before the start of each hold block. Second, we tested for treatment effects on the respiratory frequency during the paced breathing blocks. In both cases, we compared the means of these two variables across our four treatment groups by using a one-way repeated measures analysis of variance. Statistical significance was set to p<0.05 (two-tailed).

Using our first-level CVR contrasts for Hold vs Paced breathing BOLD signal changes, we conducted a set of statistical analyses to investigate the effects of task and treatment. First, to confirm that our Hold blocks, as compared to Paced breathing, elicited the expected global pattern of BOLD signal increase across the whole brain, we averaged the Hold vs Paced breathing contrast maps across treatment levels for each subject and then investigated the main effect of task by conducting a one-sample T-test at the whole-brain level. We tested the effect of task using two directed T-contrasts: one for increases (Hold>Paced) and another for decreases in the BOLD signal (Hold<Paced). Second, we investigated treatment-related effects on CVR in a one-way repeated measures analysis of variance. We tested the effect of treatment using a F-contrast to investigate changes related to treatment irrespective of direction at the whole-brain level. In all tests, we used cluster-level inference at α = 0.05 using family-wise error (FWE) correction for multiple comparisons and a cluster-forming threshold of p = 0.001 (uncorrected), according to current recommendations for the parametric analysis of BOLD fMRI data^106^.

All the analyses were conducted with the researcher unblinded regarding treatment condition. Since we used a priori and commonly accepted statistical thresholds and report all observed results at these thresholds, the risk of bias in our analyses is minimal, if not null.

## Data availability

Data can be accessed from the corresponding author upon reasonable request. A reporting summary for this Article is available as a Supplementary Information file.

## Code availability

All the code used for the analyses is openly available as part of the respective toolboxes. The matlab code to implement the spin rotations can be found in https://github.com/frantisekvasa/rotate_parcellation.

## List of Supplementary Materials

Supplementary Table 1 – Treatment, time-interval and treatment x time-interval effects on self-reported alertness, mood and anxiety.

Supplementary Figure 1 – Effects of treatment, time-interval and treatment x time-interval on self-reported alertness, mood and anxiety.

Supplementary Table 2 – Treatment, time-interval and treatment x time-interval effects on self-global cerebral blood flow.

Supplementary Figure 2 – Dose-response effects of intranasal oxytocin on global cerebral blood flow.

Supplementary Figure 3 – Main effect of time-interval on regional cerebral blood flow (rCBF) at the whole-brain level.

Supplementary Figure 4 - Dose-response effects of intranasal oxytocin on the functional connectivity of the amygdala subdivisions with the remaining regions of the brain oxytocinergic circuits.

Supplementary Figure 5 - Dose-response effects of intranasal oxytocin on the functional connectivity of the oxytocinergic network in the human brain (including amygdala’s subdivisions).

Supplementary Figure 6 – Dose-response effects of intranasal oxytocin on the respiratory belt readings at the beginning of the hold blocks of our breath-hold task.

Supplementary Figure 7 - Dose-response effects of intranasal oxytocin on respiratory frequency during the paced-breathing blocks of our breath-hold task.

Supplementary Figure 8 – Whole-brain increases in BOLD (cerebrovascular reactivity) during breath hold, as compared to paced breathing (Main effect of the task).

Supplementary Figure 9 – Expression of mRNA of oxytocin pathway genes across the main amygdala subdivisions according to the Allen Brain Atlas (ABA).

Supplementary Table 3 – Participants predictions regarding treatment allocation.

Supplementary Table 4 – Visual analog scales used to assess alertness, mood and anxiety.

Supplementary Table 5 – Number of voxels in each of the amygdala’s subdivisions regions-of-interest.

Supplementary Table 6 – List of anatomical regions-of-interest used in the group-based regional cerebral blood flow-covariance analyses of functional connectivity and respective sources.

## ACKNOWLEDGMENTS

We would like to thank all volunteers contributing data to this study.

## Author contributions

YP and DM designed the study; DM and KB collected the data; NM provided medical supervision; DM, MV and OD analyzed the data; FZ provided new analytical tools; DM and YP wrote the first draft of the paper and all co-authors provided critical revisions.

