## Supplementary materials for "“Less is more”: a dose-response account of intranasal oxytocin pharmacodynamics in the human brain"

Martins et al.

**Supplementary Table 1 – Treatment, time-interval and treatment x time-interval effects on self-reported alertness, mood and anxiety.** We investigated the effects of treatment, time-interval and treatment x time-interval on self-reports of alertness, mood and anxiety, using a linear mixed model. Statistical significance was set at *p <*0.05 (two-tailed).

| **Variable** | **Main effect of treatment** | | **Main effect of time-interval** | | **Interaction**  **Treatment x Time-interval** | |
| --- | --- | --- | --- | --- | --- | --- |
|  | **F** | **p-value** | **F** | **p-value** | **F** | **p-value** |
| **Alertness** | F(3,116.649) = 0.413 | 0.413 | F(2,140.940) = 7.829 | 0.001 | F(6,79.168) = 0.638 | 0.700 |
| **Mood** | F(3,131.340) = 0.301 | 0.825 | F(2,184.100) = 1.578 | 0.209 | F(6,87.559) = 0.260 | 1.000 |
| **Anxiety** | F(3,134.141) = 0.522 | 0.668 | F(2,180.100) = 0.595 | 0.553 | F(6,88.904) = 0.549 | 0.770 |

**Supplementary Figure 1 – Effects of treatment, time and treatment x time on alertness, mood and anxiety.** We investigated the effects of treatment, time and treatment x time on alertness, mood and anxiety ratings collected immediately before (time 1) and after (time 2) drug administration and at the end of the scanning session (time 3). Statistical significance was set at *p <*0.05 (two-tailed), after correcting for multiple comparisons using the *Sidak* correction.

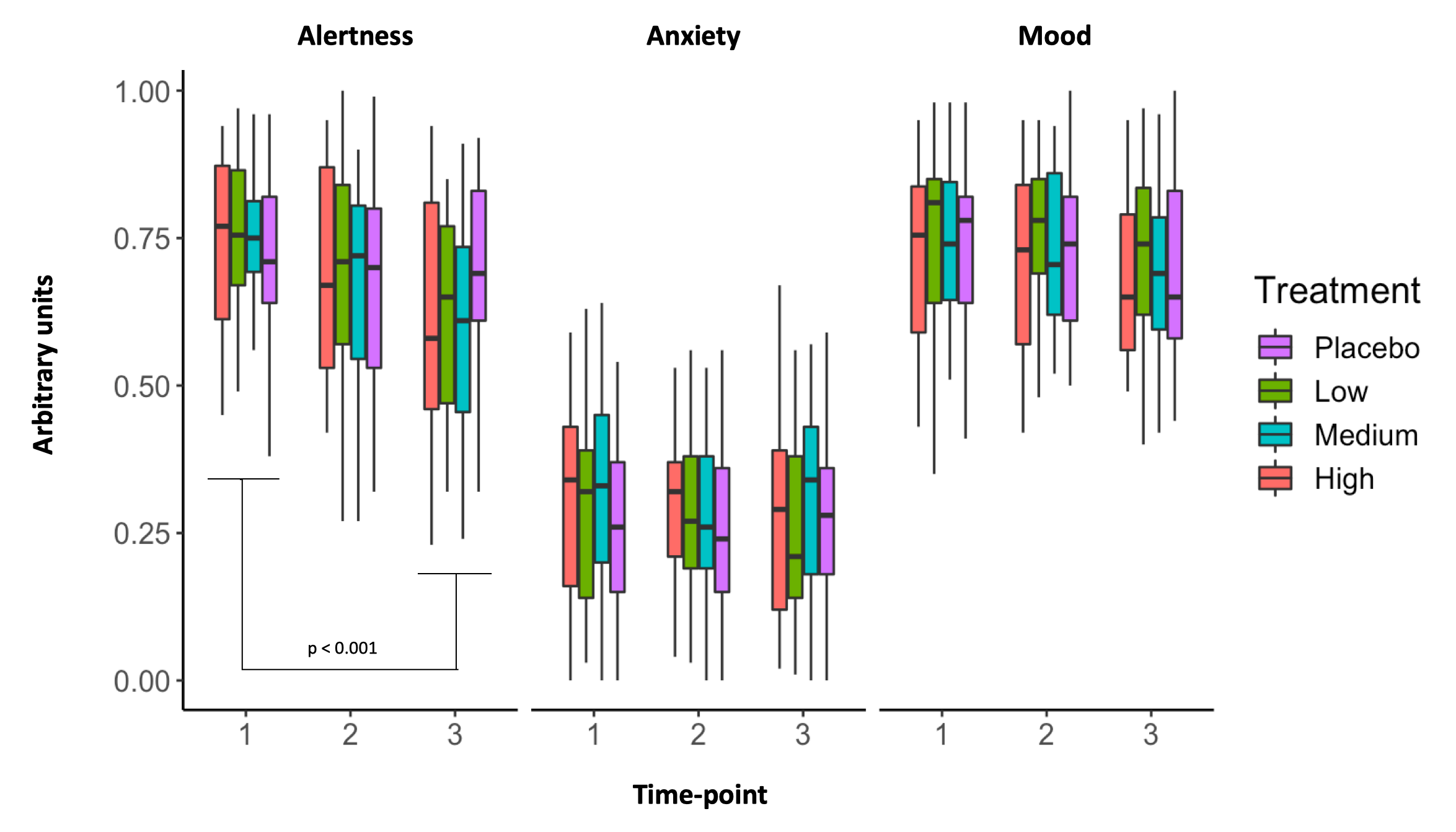

Further investigations of the significant main effect of time on alertness with direct post-hoc comparisons between time-points showed that this effect was driven by a significant decrease in the alertness ratings at time-point 3 when compared to time-point 1 (d = 0.567, p_adjusted_ < 0.001) (Supplementary Figure 1). There were no significant differences between time-points 1 and 2 (p_adjusted_ = 0.086), or time-points 2 and 3 (p_adjusted_ = 0.209).

**Supplementary Table 2 – Treatment, time-interval and treatment x time-interval effects on global cerebral blood flow (CBF).** We investigated the effects of treatment, time-interval and treatment x time-interval on global CBF, using a linear mixed model. Statistical significance was set at *p <*0.05 (two-tailed).

| **Variable** | **Main effect of treatment** | | **Main effect of time-interval** | | **Interaction**  **Treatment x Time-interval** | |
| --- | --- | --- | --- | --- | --- | --- |
|  | **F** | **p-value** | **F** | **p-value** | **F** | **p-value** |
| **Global CBF** | F(3,105.505) = 4.666 | 0.004 | (F(2,173.155) = 0.109 | 0.897 | F(6,79.441) = 0.316 | 0.927 |

**Supplementary Figure 2 – Dose-response effects of intranasal oxytocin on global cerebral blood flow (CBF).** In this graph, we present the results of the post-hoc investigations of the simple dose effects on global CBF, for which we found a significant main effect. Statistical significance was set at *p <*0.05 (two-tailed), after correcting for multiple comparisons using the *Sidak* correction.

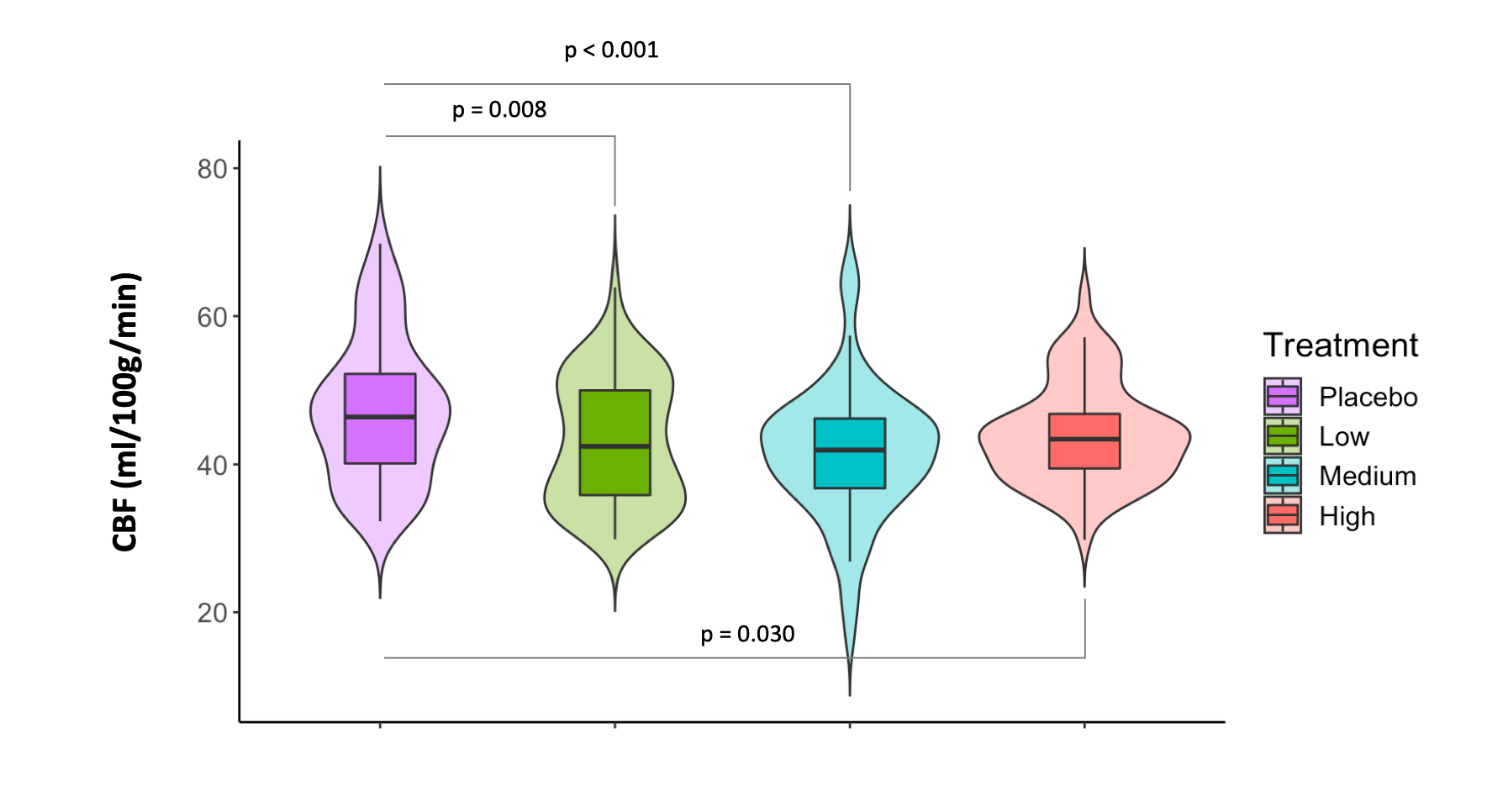

We found that all doses significantly reduced global CBF. The decrease in global CBF was maximal for the medium dose (d = 0.606, p_adjusted_ < 0.001), followed by the low dose (d = 0.456, p_adjusted_ = 0.008) and the high dose (d = 0.381, p_adjusted_ = 0.03). Follow-up comparisons between each pair of individual oxytocin doses were non-significant (smallest p_adjusted_ = 0.061).

**Supplementary Figure 3 – Main effect of time-interval on regional cerebral blood flow (rCBF) at the whole-brain level.** Clusters showing a significant main effect of time-interval interaction in rCBF, as identified in F contrasts (not capturing the direction of the change in rCBF). Global CBF was used as a nuisance variable. We conducted cluster-level inference, reporting clusters significant at p < 0.05 FWE-corrected (cluster-forming threshold: p<0.005, uncorrected). Images are shown as F-statistics in radiological convention.

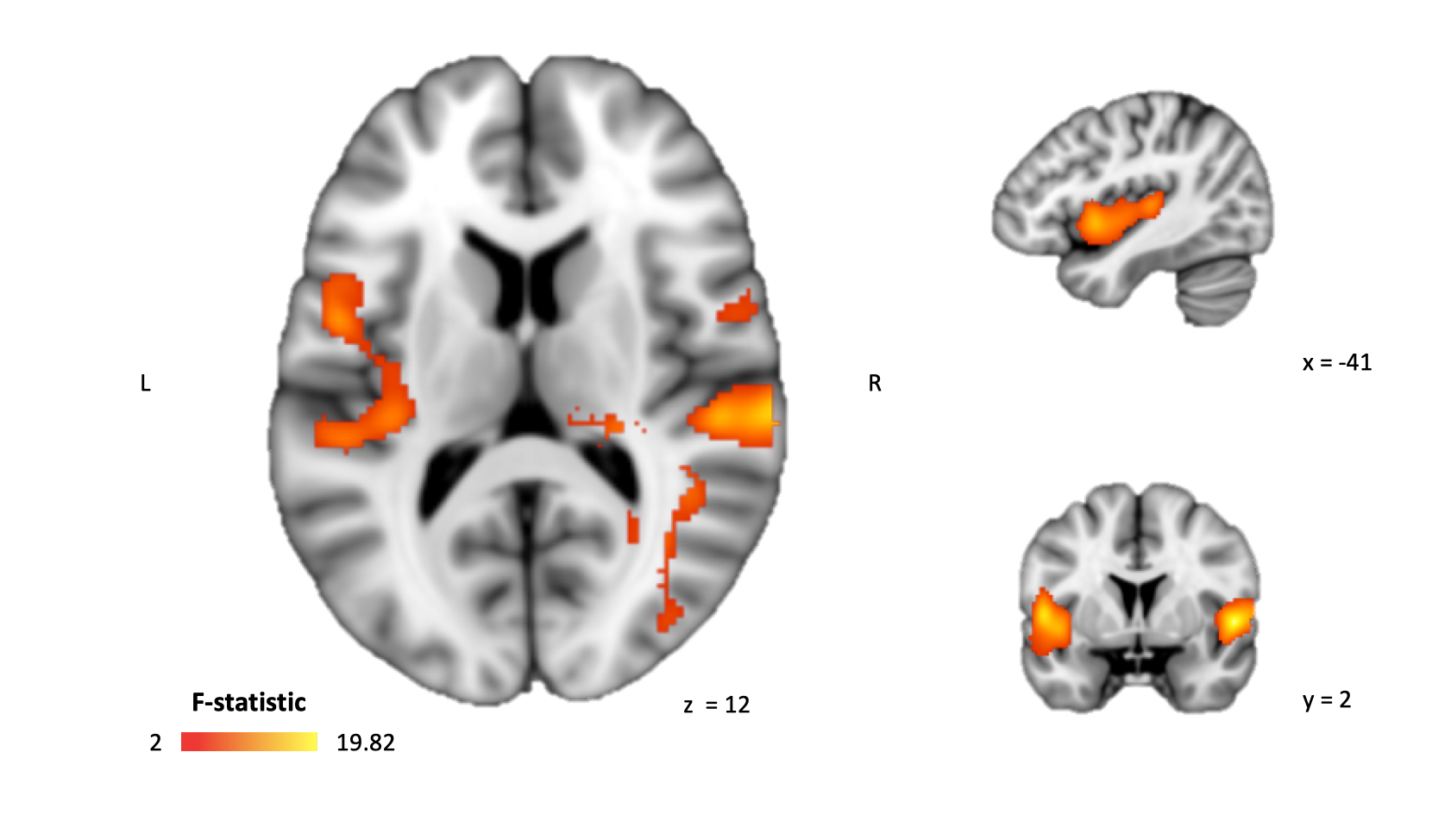

**Supplementary Figure 4 - Dose-response effects of intranasal oxytocin on the functional connectivity of the amygdala subdivisions with the remaining regions of the brain oxytocinergic circuits.** We investigated the effects of each dose of intranasal oxytocin, when compared to placebo, on the functional connectivity of each amygdala subdivision with the remaining regions-of-interest (ROI) of our brain oxytocinergic network. Instead of conducting statistical analyses on each ROI-to-ROI connections of our network, we summarized the properties of the amygdala’s connections using two graph-theory modelling metrics, node strength and clustering coefficient. Then, we compared node strength and clustering coefficient between each dose and placebo, using permutation testing (up to 10000 permutations). Statistical significance was set at *p* < 0.05 (two-tailed). CM – Centromedial; LB – Laterobasal; SF – Superficial; AStr – Amygdalostriatal transition area; R – Right; L - Left.

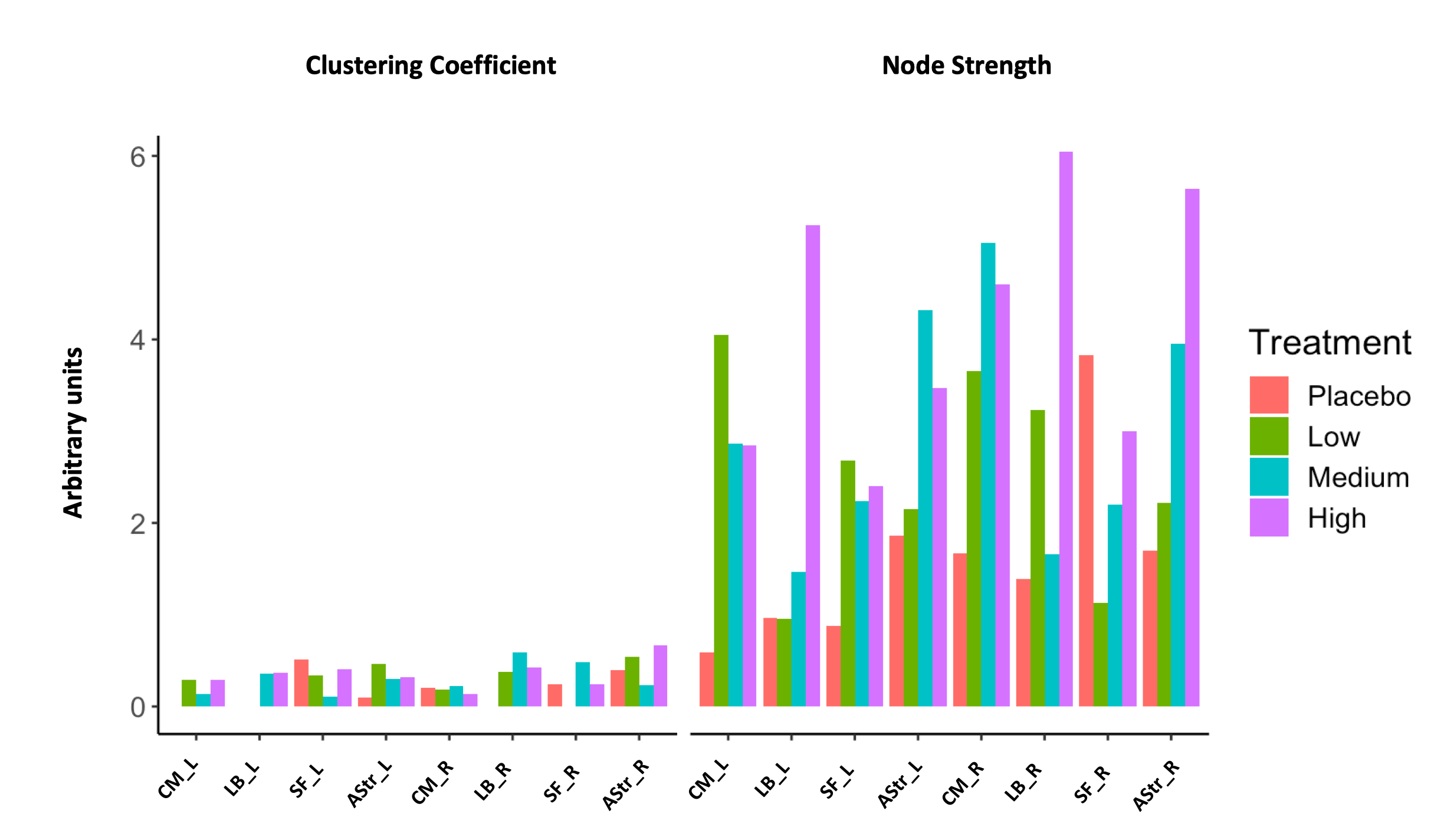

**Supplementary Figure 5 - Dose-response effects of intranasal oxytocin on the functional connectivity of the oxytocinergic network in the human brain (including amygdala’s subdivisions).** We repeated the same analysis originating Fig. 3 of the main manuscript, but this time switching the whole-amygdala regions-of-interest (ROI) by the respective centromedial, laterobasal, superficial and amygdalostriatal transition area subdivisions ROIs. Therefore, instead of four symmetric 31 x 31, we used 37 x 37 ROIs matrices (which we show for each treatment group in this figure). We then used the same approach to assess treatment related-effects using Pearson’s cross-correlations between the lower triangles of each of our three doses covariance matrices and the one from placebo (reference) as a measure of between matrices similarity. Please note that these cross-correlations were calculated using only the significant correlations (p < 0.05) present in the two matrices of each pair (i.e. elements of the matrices that overlap, after applying threshold) – we did not include all other non-significant correlations to reduce noise from potential spurious correlations. For each of these three cross-correlations we show the Pearson’s correlation coefficient *(r_pearson_)* and the correspondent 95% confidence interval and *p*-value. Statistical significance was set at *p* < 0.05 (two-tailed).

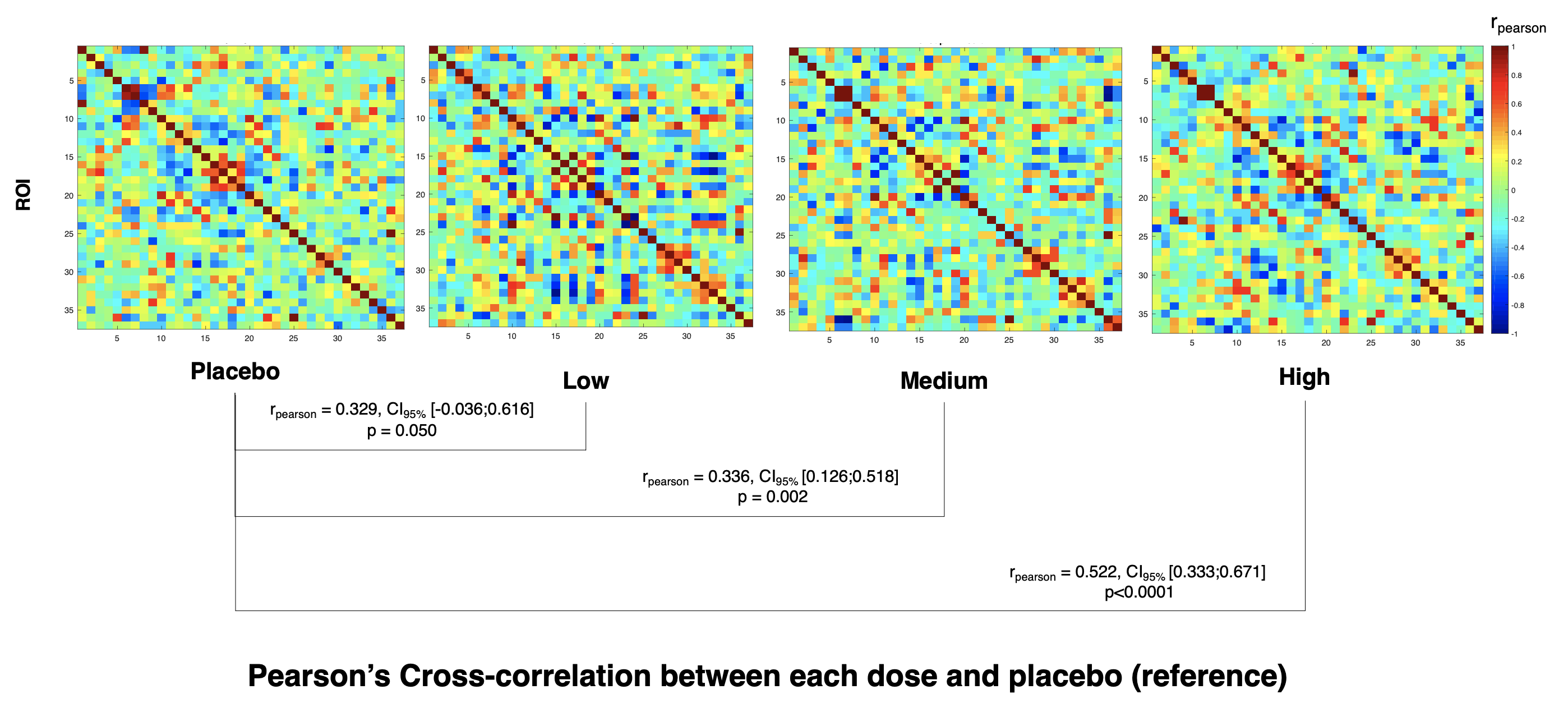

When we compared directly the cross-correlation coefficient of the associations between each dose and placebo across doses, we found that the cross-correlation coefficients for the low and medium doses were not significantly lower than the cross-correlation coefficient for the high dose but were at trend-level (low versus high: Z = -1.394, p = 0.082; medium versus high: Z = -1.395, p = 0.081). Direct comparisons of the cross-correlation coefficients between the low and medium doses yielded no significant differences (Z = -0.026, p = 0.490).

**Supplementary Figure 6 – Dose-response effects of intranasal oxytocin on the respiratory belt readings at the beginning of the hold blocks of our breath-hold task.** We investigated whether intranasal oxytocin could affect the amplitude of the last exhalation right before the beginning of the hold blocks of our breath-hold task to discard this potential confounder. For that, we retrieve the readings of the respiratory belt at the moment our participants started each hold block. First, we averaged these readings across blocks, within-each participant/session. Then, we investigated treatment effects on these mean readings using a repeated-measures one-way analyses of variance, using treatment as a factor. Statistical significance was set at *p <* 0.05 (two-tailed).

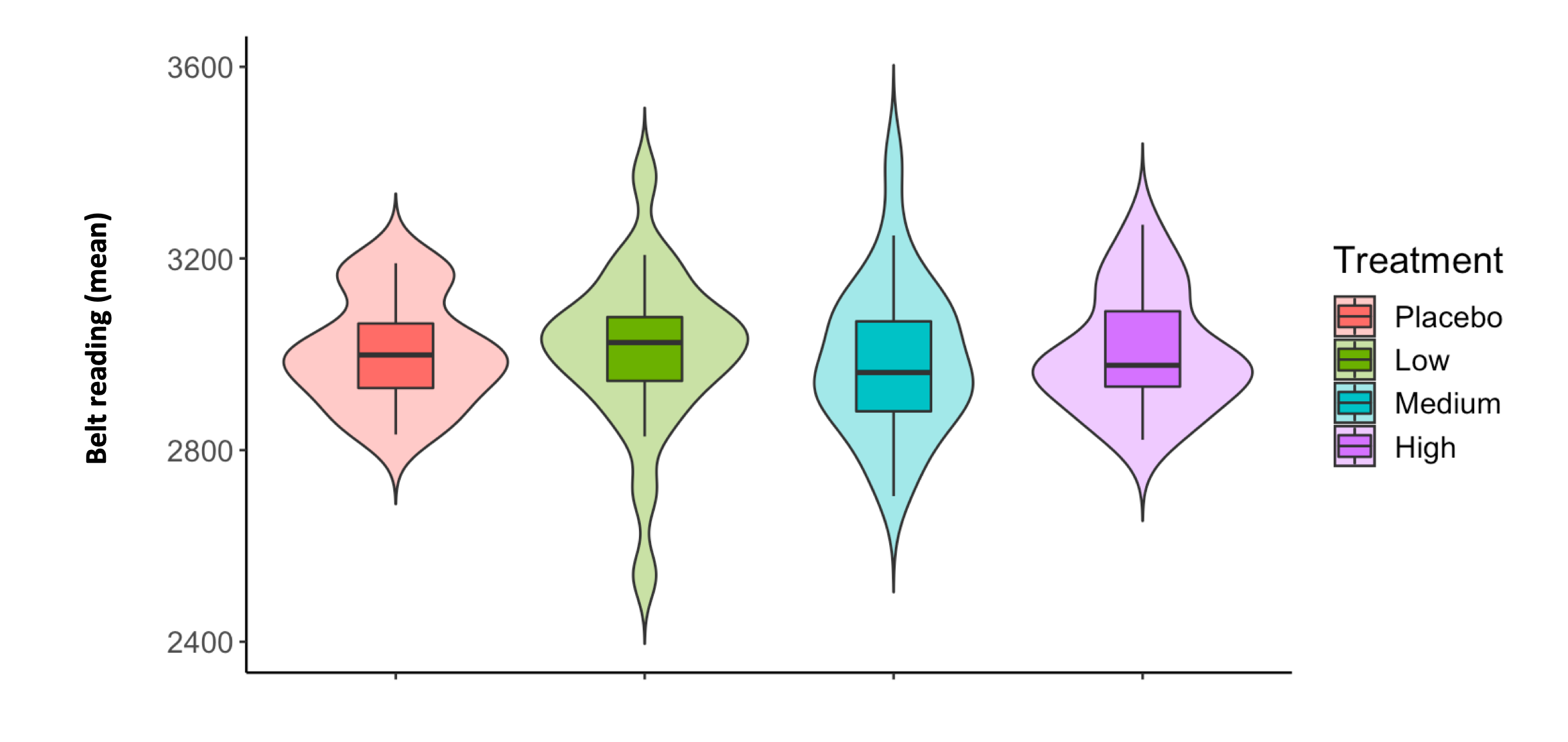

**Supplementary Figure 7 - Dose-response effects of intranasal oxytocin on respiratory frequency during the paced-breathing blocks of our breath-hold task.** We investigated whether intranasal oxytocin could affect respiratory frequency (RF) during the paced-breathing blocks of our breath-hold task to discard this potential confounder. For that, we retrieve the RF for each paced-breathing block. First, we averaged these measurements across blocks, within-each participant/session. Then, we investigated treatment effects on RF during paced-breathing using a repeated-measures one-way analyses of variance, using treatment as a factor. Statistical significance was set at *p <* 0.05 (two-tailed).

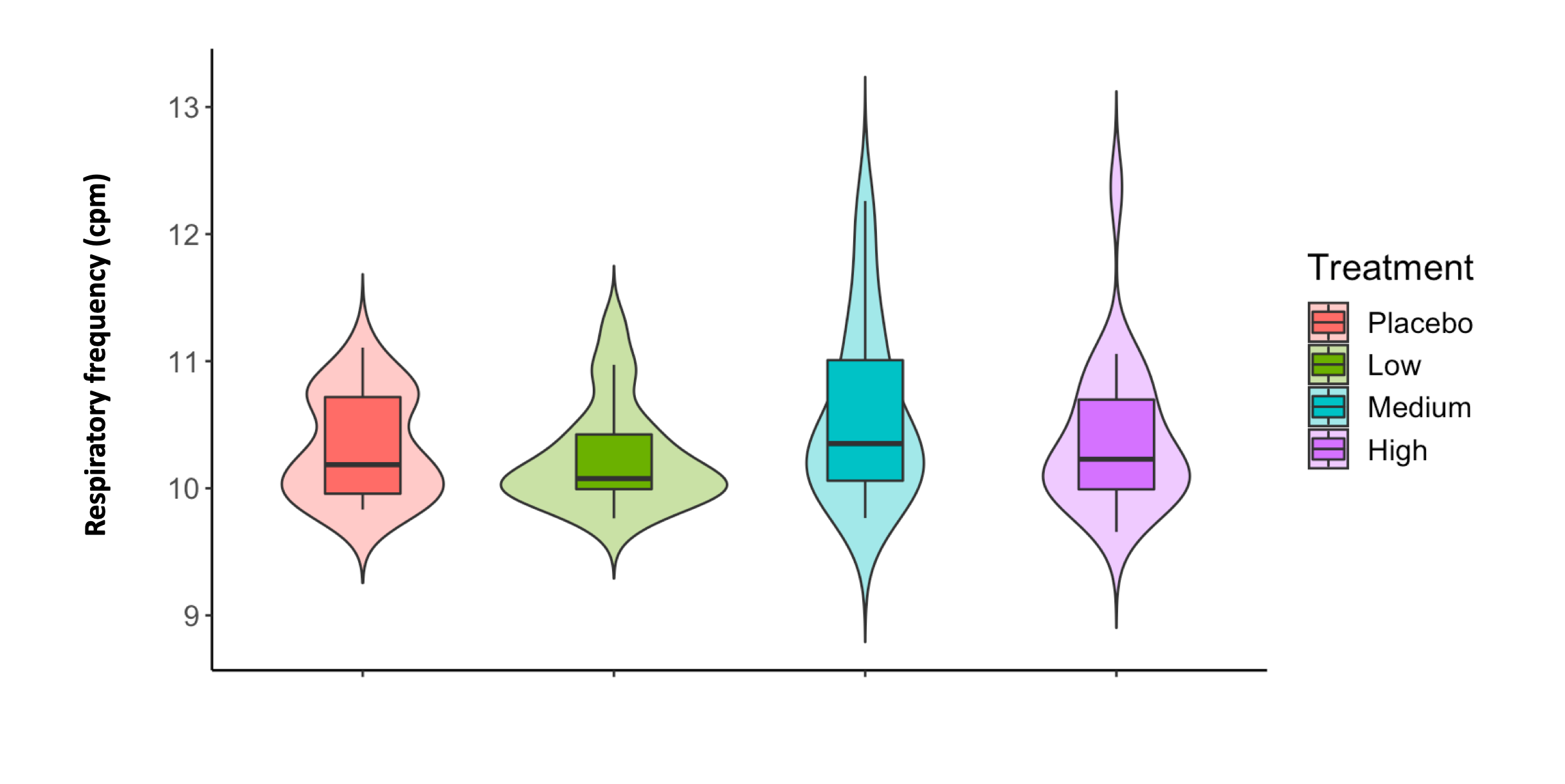

**Supplementary Figure 8 – Whole-brain increases in BOLD (cerebrovascular reactivity) during breath hold, as compared to paced breathing (Main effect of the task).** Cluster showing a significant main effect of task in BOLD signal, as identified in a T contrast capturing increases during breath hold, as compared to paced breathing (Hold > Paced). To examine this main effect of task, we pooled together the first level contrasts Hold versus Paced from all scans of our four treatment groups and conducted a one-sample T test. We conducted cluster-level inference, reporting clusters significant at p < 0.05 FWE-corrected (cluster-forming threshold: p<0.001, uncorrected). Images are shown as T-statistics in radiological convention.

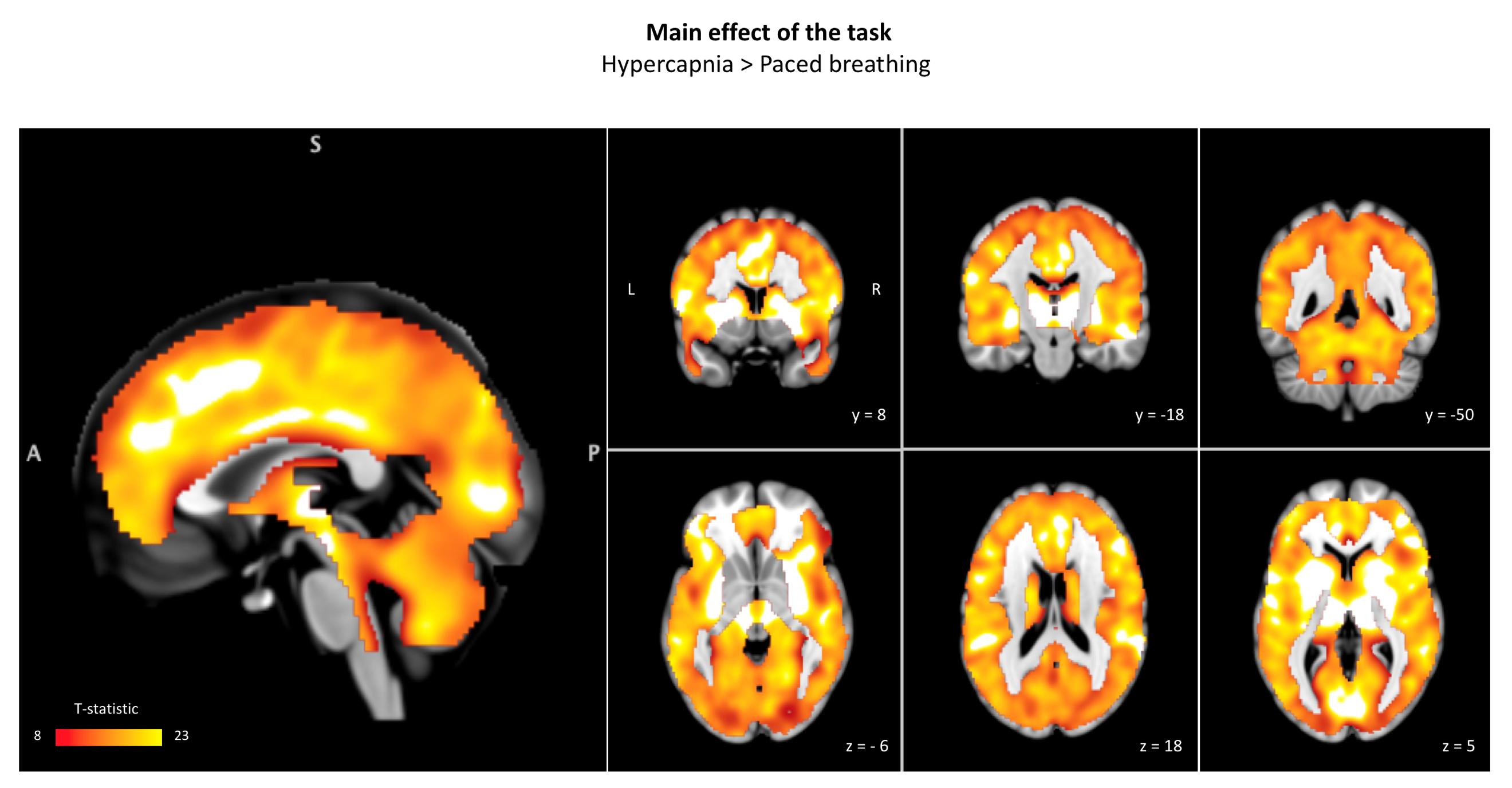

**Supplementary Figure 9 – Expression of mRNA of oxytocin targets genes across the main amygdala subdivisions according to the Allen Brain Atlas (ABA).** We retrieved from the ABA data on the level of expression of two targets oxytocin, the oxytocin receptor (OXTR) and the vasopressin receptor AVPR1A. For each donor, we matched the list of available structures in the ABA to the main amygdala subdivisions regions-of-interest used in this study. For each donor and each subdivision, we averaged the log_2_ intensity levels of mRNA expression to obtain a single measure of gene expression for that subdivision. In this figure, we present, for each of our three genes of interest, the levels of mRNA in centromedial, laterobasal and superficial subdivisions of the amygdala. We did not conduct statistical analyses for mean differences across subdivisions because the small number of donors available in the ABA would not allow us to interpret such inferences with confidence. Instead, we used this figure for illustrative purposes and to help us to interpret our findings for the effects of different doses of intranasal oxytocin on the rCBF of the different subdivisions of the amygdala.

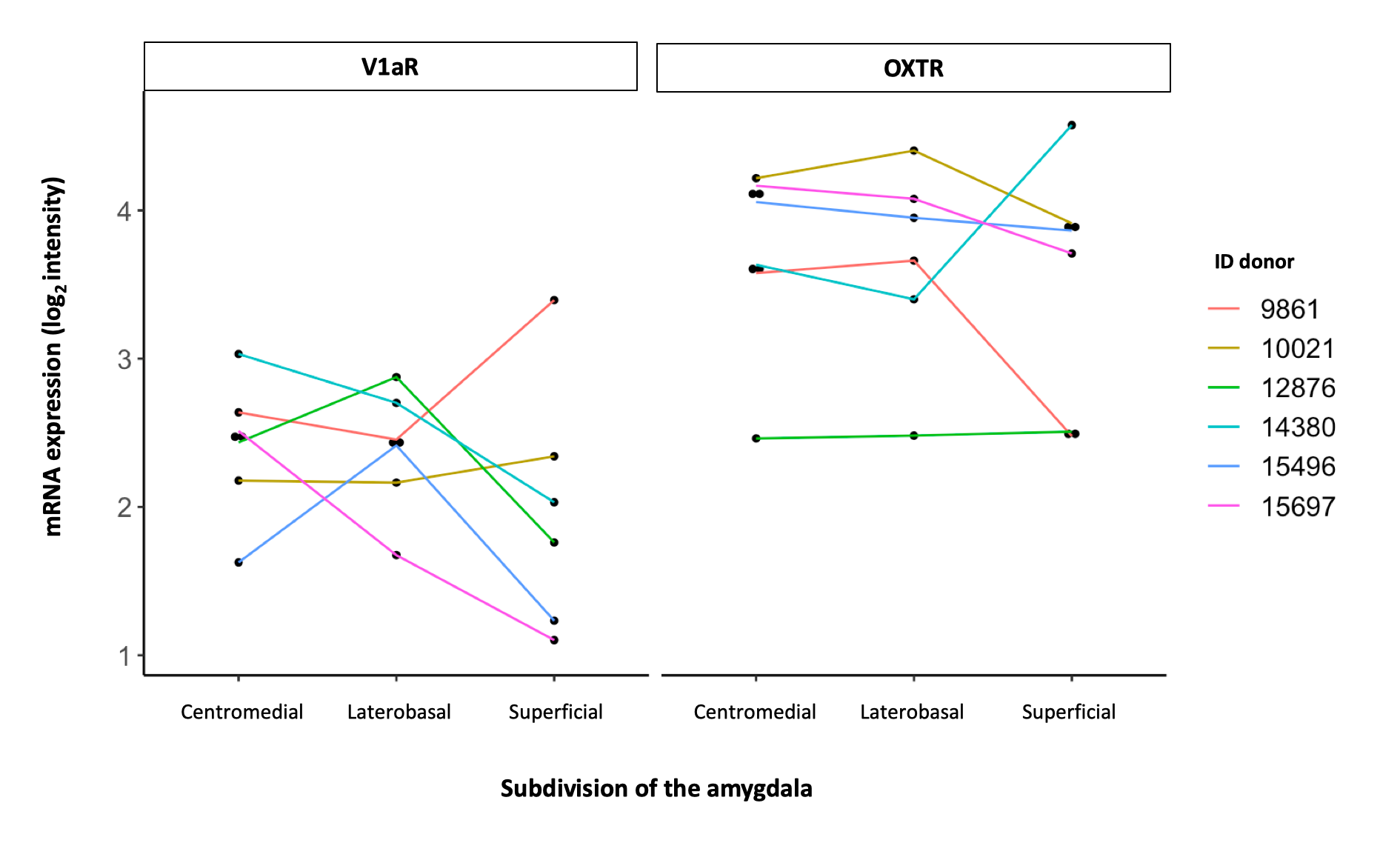

**Supplementary Table 3 – Participants predictions regarding treatment allocation.** In this cross-table, we present participants predictions regarding treatment allocation during each session.

|  | | **Participants’ prediction** | | | | |
| --- | --- | --- | --- | --- | --- | --- |
|  |  | **High** | **Medium** | **Low** | **Placebo** | **Total** |
| **True** | **High** | 1 | 5 | 6 | 12 | 24 |
|  | **Medium** | 3 | 6 | 11 | 4 | 24 |
|  | **Low** | 7 | 4 | 8 | 5 | 24 |
|  | **Placebo** | 2 | 6 | 7 | 9 | 24 |
|  | **Total** | 13 | 21 | 32 | 30 | 96 |

**Supplementary Table 4 – Visual analog scales used to assess alertness, mood and anxiety.** In this table we present the anchors for each of the 16 visual analog scales (VAS) used to assess self-reported alertness, mood and anxiety.

| **Visual Analog Scale** | **Subscale** |
| --- | --- |
| Drowsy – Alert | Alertness |
| Calm – Excited | Anxiety |
| Feeble – Strong | Alertness |
| Muzzy – Clear-headed | Alertness |
| Clumsy – Well-coordinated | Alertness |
| Lethargic - Energetic | Alertness |
| Discontented – Contented | Mood |
| Troubled - Tranquil | Anxiety |
| Mentally slow – Quick-witted | Alertness |
| Tense – Relaxed | Anxiety |
| Dreamy – Attentive | Alertness |
| Incompetent - Proficient | Alertness |
| Sad – Happy | Mood |
| Antagonistic - Amicable | Mood |
| Bored – Interested | Mood |
| Withdrawn - Gregarious | Mood |

**Supplementary Table 5 – Number of voxels in each of the amygdala’s subdivisions regions-of-interest.**

| **Region-of-interest** | **Number of voxels** |
| --- | --- |
| **Right Amygdalostriatal Transition Area** | 124 |
| **Right Centromedial amygdala** | 222 |
| **Right Laterobasal amygdala** | 1699 |
| **Right Superficial amygdala** | 387 |
| **Left Amygdalostriatal Transition Area** | 171 |
| **Left Centromedial amygdala** | 345 |
| **Left Laterobasal amygdala** | 1938 |
| **Left Superficial amygdala** | 291 |

**Supplementary Table 6 – List of anatomical regions-of-interest used in the group-based regional cerebral blood flow-covariance analyses of functional connectivity and respective sources.**

| **Anatomical region-of-interest** | **Source** |
| --- | --- |
| Right Thalamus | Harvard-Oxford Brain Atlas |
| Left Thalamus | Harvard-Oxford Brain Atlas |
| Subcallosal cortex | Harvard-Oxford Brain Atlas |
| Right Putamen | Harvard-Oxford Brain Atlas |
| Left Putamen | Harvard-Oxford Brain Atlas |
| Precuneus | Harvard-Oxford Brain Atlas |
| Medial Prefrontal cortex | Area FPm from the connectivity-based parcellation map in Franz-Xaver Neubert et al.**^1^** |
| Posterior cingulate | Harvard-Oxford Brain Atlas |
| Right Pallidum | Harvard-Oxford Brain Atlas |
| Left Pallidum | Harvard-Oxford Brain Atlas |
| Right Parahippocampal gyrus | Harvard-Oxford Brain Atlas |
| Left Parahippocampal gyrus | Harvard-Oxford Brain Atlas |
| Right Paracingulate cortex | Harvard-Oxford Brain Atlas |
| Left Paracingulate cortex | Harvard-Oxford Brain Atlas |
| Olfactory region | Automatic Anatomical Labeling atlas (version 2) |
| Right Insula | Harvard-Oxford Brain Atlas |
| Left Insula | Harvard-Oxford Brain Atlas |
| Hypothalamus | High-resolution probabilistic atlas of human subcortical brain nuclei**^2^** |
| Right Hippocampus | Harvard-Oxford Brain Atlas |
| Left Hippocampus | Harvard-Oxford Brain Atlas |
| Basal Forebrain | Anatomy toolbox |
| Right Frontal orbital cortex | Harvard-Oxford Brain Atlas |
| Left Frontal orbital cortex | Harvard-Oxford Brain Atlas |
| Right Caudate | Harvard-Oxford Brain Atlas |
| Left Caudate | Harvard-Oxford Brain Atlas |
| Brainstem | Harvard-Oxford Brain Atlas |
| Right Accumbens | Harvard-Oxford Brain Atlas |
| Left Accumbens | Harvard-Oxford Brain Atlas |
| Anterior Cingulate | Harvard-Oxford Brain Atlas |
| Right Centromedial amygdala | Anatomy toolbox |
| Right Laterobasal amygdala | Anatomy toolbox |
| Right Superficial amygdala | Anatomy toolbox |
| Right Centromedial amygdala | Anatomy toolbox |
| Right amygdala (whole) | Anatomy toolbox |
| Left Centromedial amygdala | Anatomy toolbox |
| Left Laterobasal amygdala | Anatomy toolbox |
| Left Superficial amygdala | Anatomy toolbox |
| Left Centromedial amygdala | Anatomy toolbox |
| Left amygdala (whole) | Anatomy toolbox |
